# Extracellular matrix flow guides in-vitro epithelial morphogenesis

**DOI:** 10.1101/2022.10.21.513217

**Authors:** Abdel Rahman Abdel Fattah, Francesca Sgualdino, Suresh Poovathingal, Kristofer Davie, Pieter Baatsen, Katlijn Vints, Natalia Gunko, Adrian Ranga

**Affiliations:** Laboratory of Bioengineering and Morphogenesis, Biomechanics Section, Department of Mechanical Engineering, KU Leuven, Leuven, Belgium; Center for Brain & Disease Research, VIB-KU Leuven, Leuven, Belgium; Electron Microscopy Platform of the VIB Bio Imaging Core at the KU Leuven, Leuven, Belgium

## Abstract

Changes in spatial localization of the extracellular matrix (ECM) are necessary for establishing morphogenesis in multiple developmental contexts. Both ECM motion and tissue deformation require multicellular scale coordination, however the interplay between them remain largely unexplored. Here, we reveal a novel mechanism coupling morphogenetic events and epithelia-driven ECM flow *in vitro*. We show that exposure to ECM components triggers periodic morphogenesis of dome-shaped structures in human pluripotent stem cell (hPSC) monolayers, driven by directional ECM flow. We show that this flow is initiated by local symmetry breaking events, is driven by microvilli and requires unperturbed flow conditions, microvilli function, and cytoskeletal contractility. An *in silico* model shows that a reaction-diffusion-like mechanism is responsible for organizing local morphogenesis into global tissue-wide events. We validate this model by predicting changes in cell patterning landscape during mesoderm differentiation, and demonstrate changes in cellular identity by immunohistochemistry and scRNAseq. These results demonstrate that transport of ECM over epithelia, termed ECM flow, is a major contributor in sustaining morphogenesis and differentiation. Our *in vitro* approach suggests that ECM flow may be a broadly conserved mechanism guiding multi-cellular morphogenesis and may be further explored to investigate the role of ECM transport in other model systems.

## Introduction

During embryonic development, epithelia achieve their form through coordinated changes in cell identity and position in space. Initial symmetry-breaking events in homogenous tissues trigger collective cellular dynamics which follow fluid-like behaviors guided by mechanical forces^1–3^. Much like cellular flows, the extracellular matrix (ECM), which is in constant engagement with morphing tissues, also undergoes important but much less explored coordinated fluid-like motion^4^. At the earliest stages of development, ECM components are trafficked between the hypoblast and epiblast to signal downstream developmental processes^5^. At the onset of gastrulation, convergent and divergent points help establish convective-like ECM currents that define and guide primitive streak formation^6^. Subsequently, ECM flows during axis elongation help direct cellular streams and modulate cell mobility, velocity and arrangement in the caudal presomitic mesoderm^7^. The dynamic nature of the ECM across all stages of development raises the question as to the link between ECM flow and morphogenesis, and to the driving force behind ECM mobility, which has remained largely underexplored^4^.

*In vitro* systems that combine ECM components and pluripotent stem cell-derived multi-cellular constructs offer promising experimental setups to advance our understanding of the interplay between ECM and morphogenesis. The ECM can act to impose mechanical constrains on embedded tissues and subject them to mechanical forces. For example, increasing the stiffness of polyethylene glycol (PEG) hydrogels reduces the growth rate of embedded human neural tube organoids. However, ECM components can also act a signaling factors that regulate cellular processes. For example, the presence of laminin in PEG hydrogels increases dorsoventral patterning and proliferation of embedded mouse neural tube organoids. More recently, the exogeneous effects of the ECM have been demonstrated in impressive embryo-like structures. For example, exposure to Matrigel, a naturally derived ECM, resulted in the emergence of *in vivo*-like neural tube and somite structures^10^ together with axis elongation in mouse^11^ and human^12^ gastruloids, as well as neural tube folding using micropatterned hPSCs^13^. The underlying mechanisms of these cell-ECM interactions, however, is not well understood. A reciprocal relation between ECM and tissue morphogenesis may exist, as has been recently demonstrated *ex vivo* using avian dermal cells that interact with a collagen microenvironment^14^. However, the study focused on mesenchymal cells where ECM components did not flow independently of cells as demonstrated in other systems^6,7^. Thus whether feedback in ECM-tissue interplay is required for epithelial morphogenesis remains to be elucidated. As such, we lack a fundamental understanding of the role of ECM flow during epithelial morphogenesis, particularly during critical symmetry-breaking events in the early embryo.

Here, we demonstrate spontaneous and periodic morphogenetic events in hPSC monolayers upon exposure to Matrigel, and show that this phenomenon requires ECM flow in the direction of the symmetry-breaking event. The development of an *in silico* model based on these experimental observations allowed us to recapitulate and predict *in vitro* outcomes. We employ single cell RNA sequencing to reveal differentially expressed genes and processes between hPSC monolayer exposed to Matrigel and those in control conditions. By investigating the apical cell surface using time-lapse and scanning electron microscopy, we examine the role of apical microvilli motion in driving ECM flow. Finally, we explore the role of ECM flow in guiding morphogenesis while defining the cell patterning landscape. Our model system establishes the requirement for feedback between ECM flow and multi-epithelial cellular collectives in the establishment and maintenance of morphogenesis.

## Symmetry-breaking events trigger directed ECM flow

In order to investigate the effect of ECM on epithelial morphogenesis, we first developed a minimal in-vitro model system where we supply Matrigel diluted in growth medium to confluent hPSC monolayers. Growth medium supplemented with 10% Matrigel produced cellular dome structures with regular periodicity (156.2 ± 42.3 μm) across the monolayer, compared to Matrigel-free control conditions, which remained unpatterned (**Fig.1a**). This characteristic morphogenetic response was accompanied by high accumulations of laminin, a major component of Matrigel, having a toroidal shape above the domes (**Fig.1b**). Laminin receptor 67LR expression was also highest in the domes, suggesting an active cellular response to emerging ECM toroids (**Supplementary Fig.1)**. Moreover, since Matrigel was homogenously introduced to the culture, laminin expression in Matrigel conditions suggests that active ECM activity is responsible for Matrigel toroid generation. Indeed, by fluorescently tagging exogenous laminin, we observed the redistribution and accumulation of laminin into pockets above domes (**Supplementary Fig.2**, **Supplementary Movie 1**).

**Figure 1.**
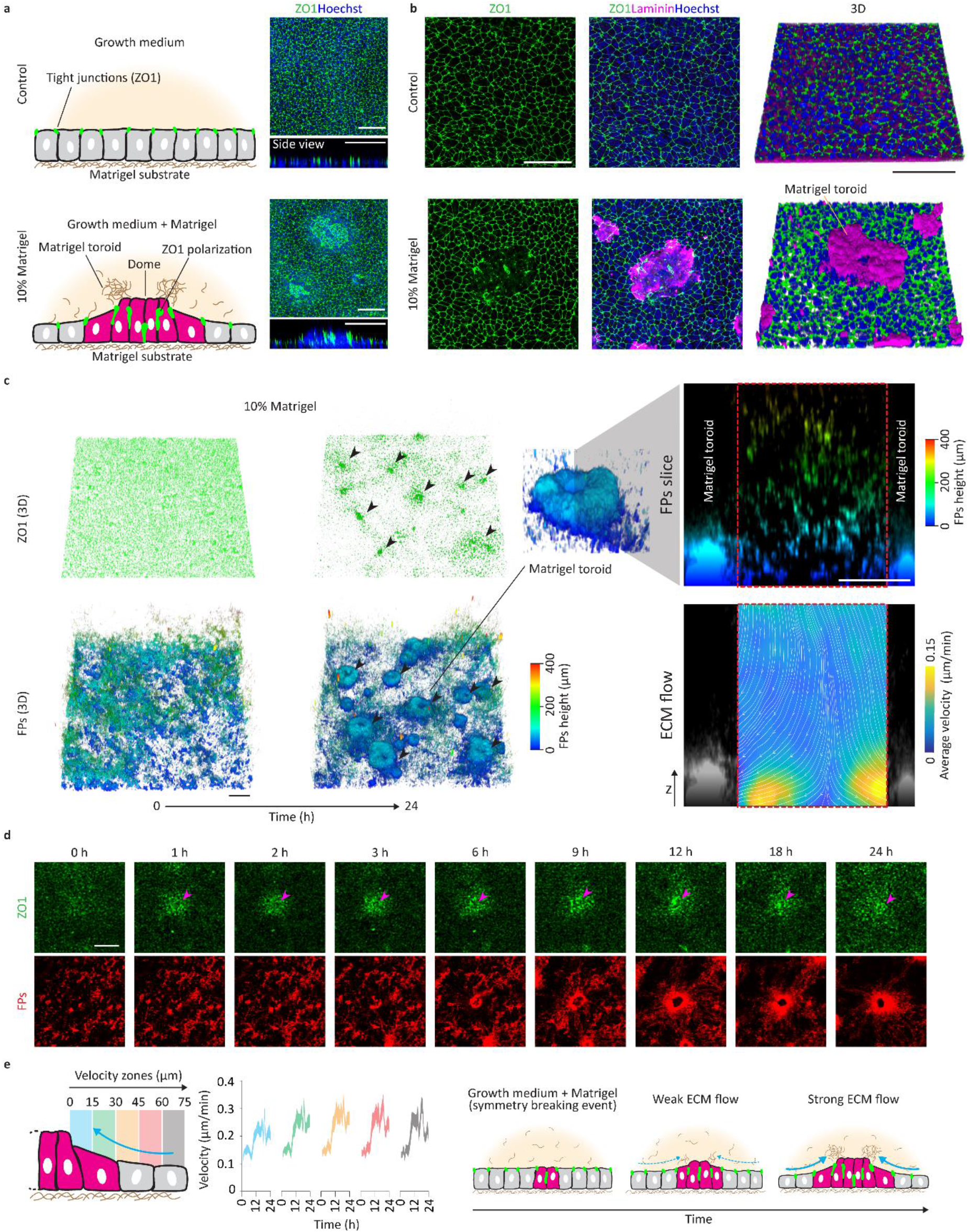
Concomitant morphogenesis and ECM flow dynamics. **a** Side and top view schematic representation of hPSC monolayers exposed to Matrigel and Matrigel-free pluripotent media. Side and top view representative images of control and Matrigel conditions at 24 h showing nuclei and ZO1 expression (n = 5). Scalebar 100 μm. **b** Representative images and 3-D reconstruction images of ZO1 and laminin expressions in 10% Matrigel and control conditions (n = 4, 10% 3-D reconstruction image rotated from top view image). **c** Representative 3-D reconstructed temporal dynamics of ZO1 expression and Matrigel-tagged FPs over 24 h. FPs channel color coded for height. Black arrow point to representative dome formation locations at 24 h in 3-D ZO1 and FPs channel. Scalebar 100 μm. Representative 3-D reconstructed image of Matrigel toroid and y-z section of FPs between two adjacent Matrigel toroids color coded for height and accompanied 24 h average velocity map color coded for velocity magnitude where white lines and arrowhead denote flow streamlines and direction. (n = 4). Scalebar 50 μm. **d** Representative images showing the temporal dynamics of a representative symmetry breaking event (ZO1 expression) and accompanied Matrigel-tagged FPs motion over 24 h at the Matrigel-monolayer interface (n = 4). Scalebars 50 μm. **e** Side view schematic representation of dome, surrounding monolayer and ECM flow velocity investigation zone. Instantaneous velocity profiles for each zone over 24 h color coded for velocity zones. Curve bounds represent SEM (n = 3 for a total of 40 domes per velocity zone). Schematic representation showing the temporal dynamics of a symmetry breaking event (ZO1 expression) and accompanied Matrigel-tagged FPs motion.

To enable a better visualization and analysis of ECM activity we tagged ECM components with fluorescent particles (FPs) and observed their time-resolved activity in relation to a monolayer of tight junction reporter (ZO1-GFP) hPSCs by live confocal microscopy. Over 24 h, the initially homogenous FP distribution gradually accumulated into toroids that corresponded to locations of high ZO1 expression (**Fig.1c**, **Supplementary Movie 2**) and continued for an additional 24 h (**Supplementary Fig.3**). Analysis by particle image velocimetry (PIV) revealed that FP accumulation was the result of Matrigel flow vectors directed towards dome positions, which represent flow convergence points (**Fig.1c**, **Supplementary Movie 3**). By contrast, between domes, divergence points exist where Matrigel flow vectors point radially outwards. We observed no lateral flow patterns in cell-free environments (**Supplementary Fig.4**, **Supplementary Movie 3**), confirming that ECM flow is a cell-driven phenomenon and occurs at the cell-Matrigel interface.

We hypothesized that ECM flow was triggered by cellular symmetry-breaking events in the initially homogeneous epithelial monolayer. Indeed, we found that every ECM-rich toroid was associated with an initial symmetry-breaking event characterized by the formation of a high ZO1 polarization (**Fig.1d**). Time-lapse imaging allowed us to determine that ZO1 polarization always occurred before ECM flow was initiated. Additionally, by quantifying flow velocities in concentric zones around each dome, initially slow ECM flow was observed, followed by accelerating velocities (up to 0.3 μm/min) between 6 and 12 h after Matrigel introduction (**Fig.1e**). We observed large variations in the magnitude of instantaneous velocity vectors at various timepoint, suggesting that ECM flow patterns are not constant (**Supplementary Fig.5**). However, flow patterns retain a consistent direction over time that permit the eventual accumulation of ECM toroids near the morphing domes.

## Local polarity guides ECM flow

We hypothesized that the observed bulk ECM motion is the result local ECM flow patterns at the cell-Matrigel interface. We therefore introduced coated iron oxide nanoparticles with FPs into our culture system, which allowed us to interrogate locally, within the vicinity of a single ECM component aggregate, ECM component-cell interaction at the cell surface **(Fig.2a)**. Upon Matrigel addition, fluorescent clusters (FCs) converged towards the domes in linear trajectories, while in the absence of Matrigel and without ECM flow, the trajectories of FCs were circular and random (**Fig.2a, Supplementary Fig.6**).

**Figure 2.**
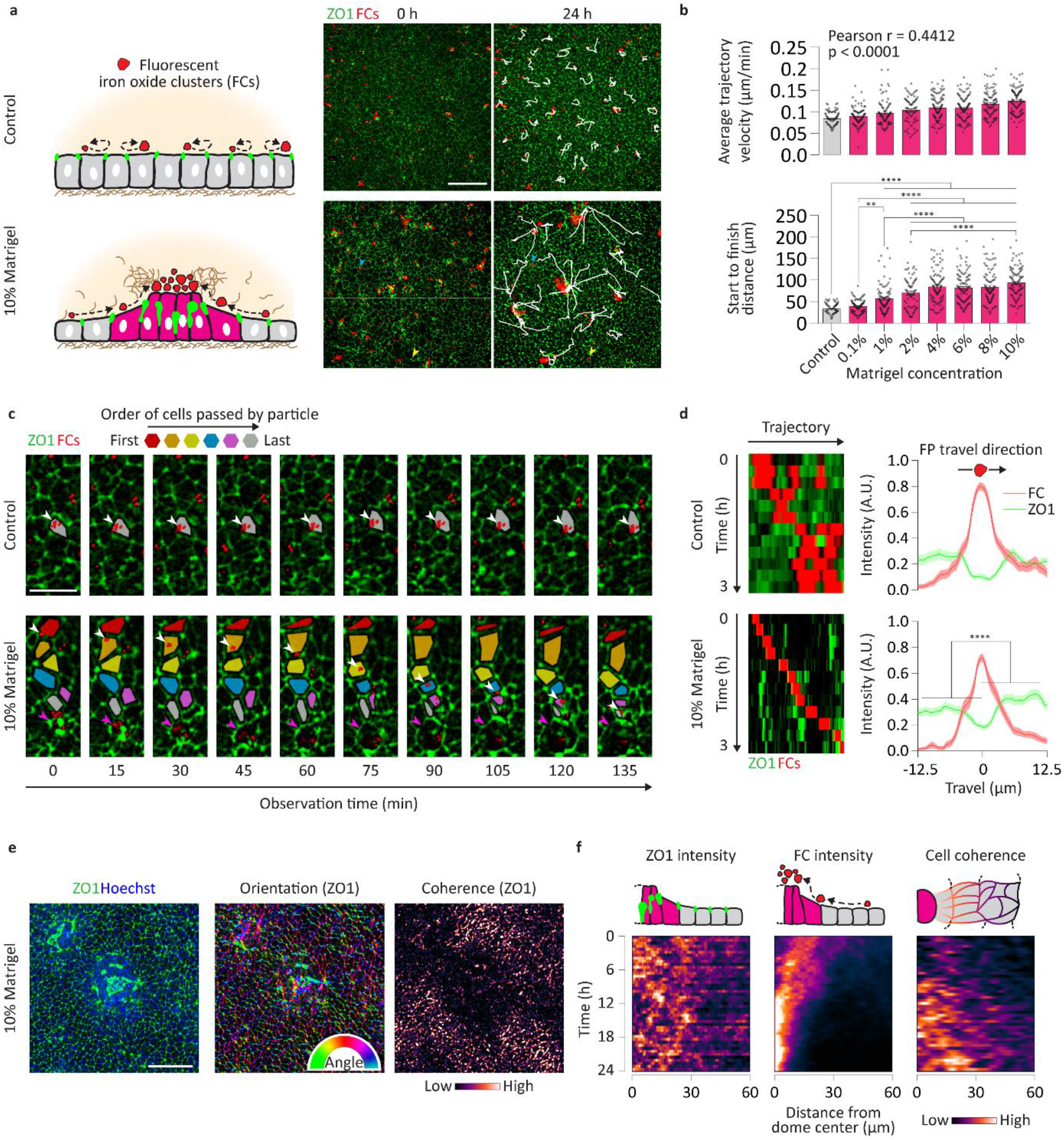
Symmetry breaking events triggers ECM flow. **a** Side view schematic presentation of control and 10% Matrigel conditions with FCs. Representative images of ZO1 expression and FC temporal dynamics with FC trajectories (white segments) after 24 h in culture. Blue and yellow arrow heads point to representative stationary FCs in 10% Matrigel conditions. Scalebar 100 μm. **b** Quantification of average trajectory velocity and trajectory start to finish distance at various Matrigel concentrations and control conditions (n = 3 for >70 trajectories per condition), *p*-values denote Pearson correlation analysis average trajectory velocity *p* < 0.0001. Statistical analysis was determined for the start to finish distance by one-way ANOVA for multiple comparisons control-1 to 10% *p* < 0.0001, 0.1-1% *p* = 0.0063, 0.1-2 to 10% *p* < 0.0001, 1-2 to 10% *p* < 0.0001, 2-10% *p* < 0.0001. Error bars are SEM. **c** Representative images of ZO1 expression and single FC (white arrowhead) dynamics in control and 10% Matrigel conditions over 135 min. Purple arrowheads depict growing symmetry breaking event in Matrigel condition. Cell shape color depict cells traversed by the FC and are color coded by order of crossing (n = 3). Scalebar 25 μm. **d** ZO1 expression and FC intensity kymographs of FC trajectories in control and 10% Matrigel conditions (n = 3 for 5 trajectories per condition). ZO1 expression and FC intensity profiles (n = 3 for 10 measurements per trajectory for a total of 50 measurements). Statistical analysis for Matrigel conditions was determined using unpaired two-sided t-test with *p* < 0.0001. Error bands are SEM. **e** Representative images of ZO1 expression, ZO1 expression orientation color coded by angle and ZO1 expression directional coherence color coded by coherence magnitude (n = 3). Scalebar 50 μm. **f** Representative images showing ZO1 expression, FC location and ZO1 expression directional coherence temporal dynamics (n = 3). Scalebar 50 μm. **c** Side view schematic representations showing ZO1 expression, FC location and ZO1 expression directional coherence. Top view schematic representation of FC location and cell geometry. Kymographs describing ZO1 expression, FC location and ZO1 average expression directional coherence dynamics color coded for magnitude (from **e**, n = 3 for a total of 30 domes).

We next assessed whether Matrigel concentration could affect ECM interactions and found that dome morphogenesis and FC accumulation could be confirmed for Matrigel concentrations >0.1% (**Supplementary Fig.7**). Additionally, Matrigel concentrations >1% did not significantly affect dome periodicity (**Supplementary Fig.8**), dome height (**Supplementary Fig.9**), FC trajectory length, or start to finish distances (**Fig.2b**).

We next investigated whether ECM flow was the result of cell motion by following individual particles, and found that while FCs remained on their host cells in control conditions, they freely traversed across multiple cells as they headed toward growing symmetry breaking events in Matrigel conditions. In this case, cell motion had little effect on trajectory direction (**Fig.2c**). By analyzing ZO1 and FC kymographs along FC trajectories, we found that local FC travel direction was associated with higher upstream ZO1 signals compared to control conditions (**Fig.2d**). This suggests that particle motion, and by extension ECM flow, are linked to local cell polarization along the monolayer plane.

Because polarity events often involve geometric rearrangement on a multicellular level, we investigated whether tissue organization and cellular alignment guided particle trajectory and ECM flow. We therefore attributed cellular alignment, based on ZO1 signal orientation, a coherence score which gradually increased (**Supplementary Fig.10**) to reach a maximum close to dome positions at 24 h after Matrigel introduction (**Fig.2e**). To further explore this correlation, we evaluated ZO1, FC and coherence kymographs (**Fig.2f**) and found that ZO1 polarization was present at the start of the symmetry breaking event, and was maintained thereafter. FC intensity dynamics indicated a transition from slow (at time < 6 h) to fast (at time > 12 h) velocities. Interestingly, this transition correlated with the emergence of established tissue coherence surrounding the domes (at time > 10 h), suggesting that tissue-level organization accelerated ECM flow. Indeed, divergent streamline points where ECM flow was weak and directionless were observed between adjacent domes where the tissue was less coherent, resulting in stationary particles in these regions (**Fig.2a** blue and yellow arrowheads). By extension, cells in these regions not directly involved in morphogenesis were not participating in directed ECM activities.

Overall, theses results suggested that changes in local polarity trigger and guide ECM flow, resulting in global morphogenetic activities and patterned ECM accumulation sites.

## ECM flow and morphogenesis interplay

In order to determine the relation between ECM flow and morphogenesis, we interrogated the spatial variation of the velocity field **V** and found its divergence ∇∙**V** to anticorrelate with dome positions, i.e. regions with high ECM accumulation and polarity, as measure by ZO1 expression (**Fig.3a**). Knowing that polarity guides ECM flow, in light of the anticorrelation (**Fig.3a**), we explored whether an established ECM flow was in turn necessary to maintain high ZO1 expressions, i.e. whether whether ECM flow acts as a positive feedback loop for morphogenesis.

**Figure 3.**
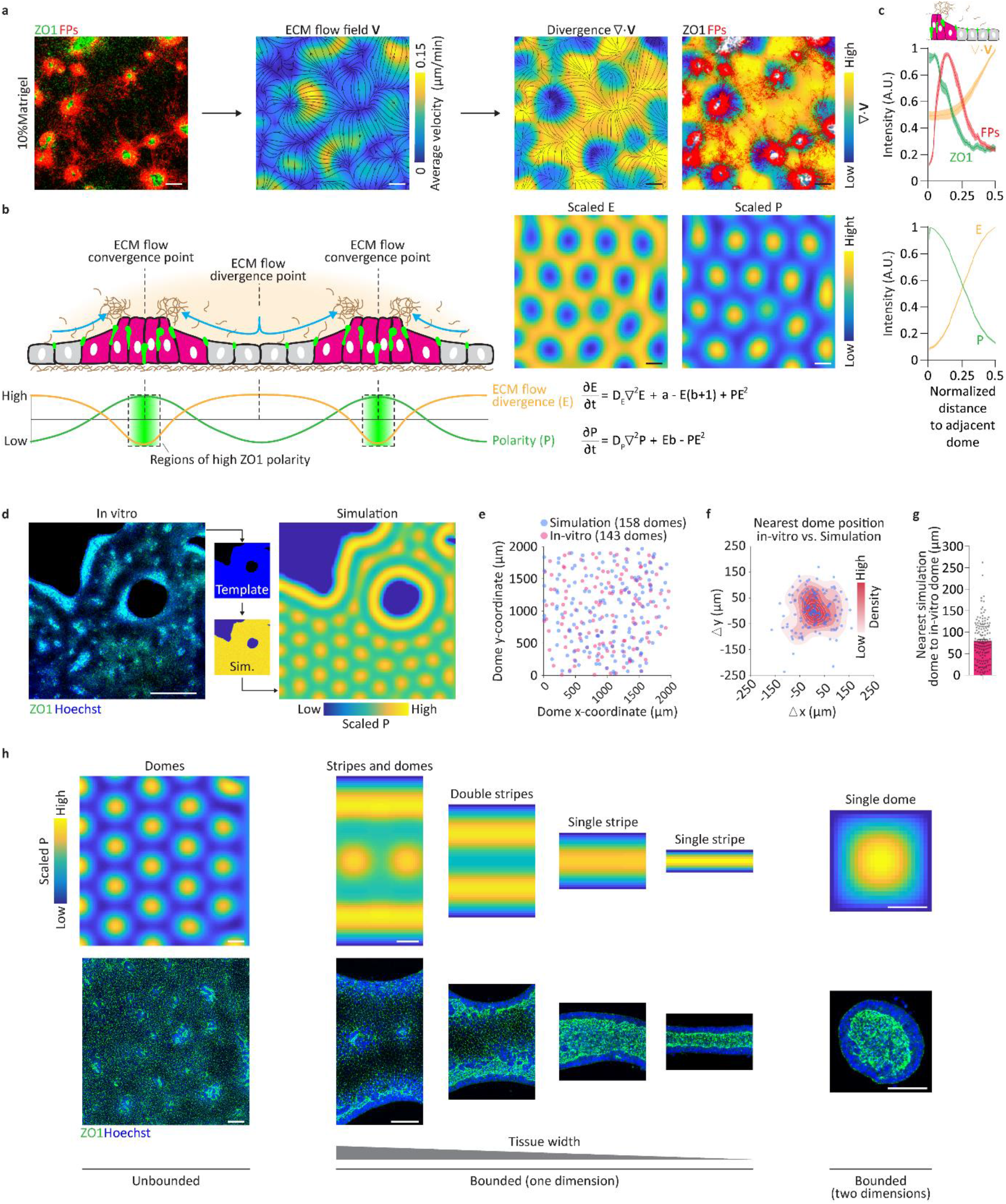
Dome morphogenesis and ECM flow anticorrelation. **a** Representative images of ZO1 expression and FP distribution at 24 h in 10% Matrigel conditions (n = 4) and corresponding 24 h average velocity map color coded for velocity magnitude where black lines and arrowhead denote flow streamlines and direction, derived divergence field color coded for divergence magnitude with superimposed streamlines, and divergence field with superimposed ZO1 expression (white) and FPs distribution (n = 3). Scalebars 100 μm. **b** Schematic representation of dome morphogenesis and ECM flow relation. Schematic profile representation of ECM flow divergence (E), tissue polarity (P) and respective mathematical expressions. Representative 2-D simulation results of scaled E and P expressions color coded for magnitude. Error bands are SEM. Scalebars 100 μm. **c** Normalized radial profiles over half the distance between two adjacent domes of ZO1 expression, FP distribution and flow divergence magnitudes (n = 3, for a total of 50 domes). Normalized radial profiles over half the distance between two adjacent *in silico* domes (defined by scaled P expressions) of scaled E and scaled P magnitudes (n = 3, for a total of 50 domes). **d** Process of converting *in vitro* tissue layout to a simulation template. Representative comparison between simulation and *in vitro* results depicting scaled P result. Scalebars 500 μm. **e** Pooled x and y coordinates of *in vitro* and *in silico* domes (n = 4 for 143 *in vitro* and 158 *in silico* domes). **f** Difference in x and y coordinates between each *in vitro* dome and the respective nearest *in silico* one (from **f**, n = 4 for 143 *in vitro* domes). **g** Average separation length between nearest *in silico* and *in vitro* domes (from **f**). Error bars are SEM. **h** Scaled expression of T for various domain spaces color coded for T signal concentration with representative images of *in vitro* similar results with comparable dimensions (n = 5). Scalebars 100 μm.

To explore this relationship further, we developed a minimal *in silico* model system based on 2-D reaction-diffusion-like principles (**Fig.3b, see methods**). Here, ZO1 polarity and ECM flow divergence are phenomenologically linked to signals P and E respectively. The signals can interact through a system of equations, which enforces a feedback between them, rendering their expressions interdependent (**Fig.3b**). In addition, the signals anticorrelate in intensities to reflect the relation between ZO1 polarity and ECM flow divergence. Therefore, regions of high P and low E expressions are *in silico* analogues of dome positions where polarity is high and the ECM accumulates. We found that the general phenotypes and radial profiles of P and E reflect those of ZO1 and ∇∙**V** respectively observed *in vitro* (**Fig.3b,c**). To quantify the model accuracy, we evaluated *in silico* dome positions produced in domains with comparable geometries to *in vitro* tissues (**Fig.3d, Supplementary Movie 5**). We found that when in-silico and *in vitro* spaces (**Fig.3e**) were superimposed, *in silico* domes were closest to their *in vitro* counterparts of similar x-coordinate ranks (**Supplementary Fig.11a, 11b)** with nearest *in silico*-*in vitro* neighbors on average ~75 μm apart (**Fig.3f,g**). This *in silico* framework also provided a powerful predictive tool (**Fig.3h**), not only recapitulating periodic dome formations in unbounded tissues, but equally predicting high polarity regions of various forms such as stripes and domes, depending on tissues boundaries and geometries. Overall, the *in silico* model results suggest a relationship between ECM-flow and morphogenesis involving feedback. To further test this hypothesis we performed *in silico* parameter perturbations targeting signals P and E and validated the results *in vitro*.

Minimizing the propagation of the polarity signal P throughout the tissue resulted in constantly high P values, indicating that the entire domain becomes a single region of high ZO1 polarization and constant low E values, suggesting that ECM flow was abrogated throughout (**Fig.4a**). As predicted by the *in silico* model, perturbation of the in-vitro model with Blebbistatin, a known inhibitor of contractility and modulator of cell polarity, we found high ZO1 polarization into lumen-like structure throughout the tissue, absence of the monolayer phenotype, and no evidence of ECM-flow (**Fig.4a**). Interestingly, the model predicted dome morphogenesis to be highly sensitive to polarity perturbations, where their abrogation can be achieved by weak polarity impairment, a trend reflected *in vitro* through very low concentrations of Blebbistatin (**Supplementary Fig.12**).

**Figure 4.**
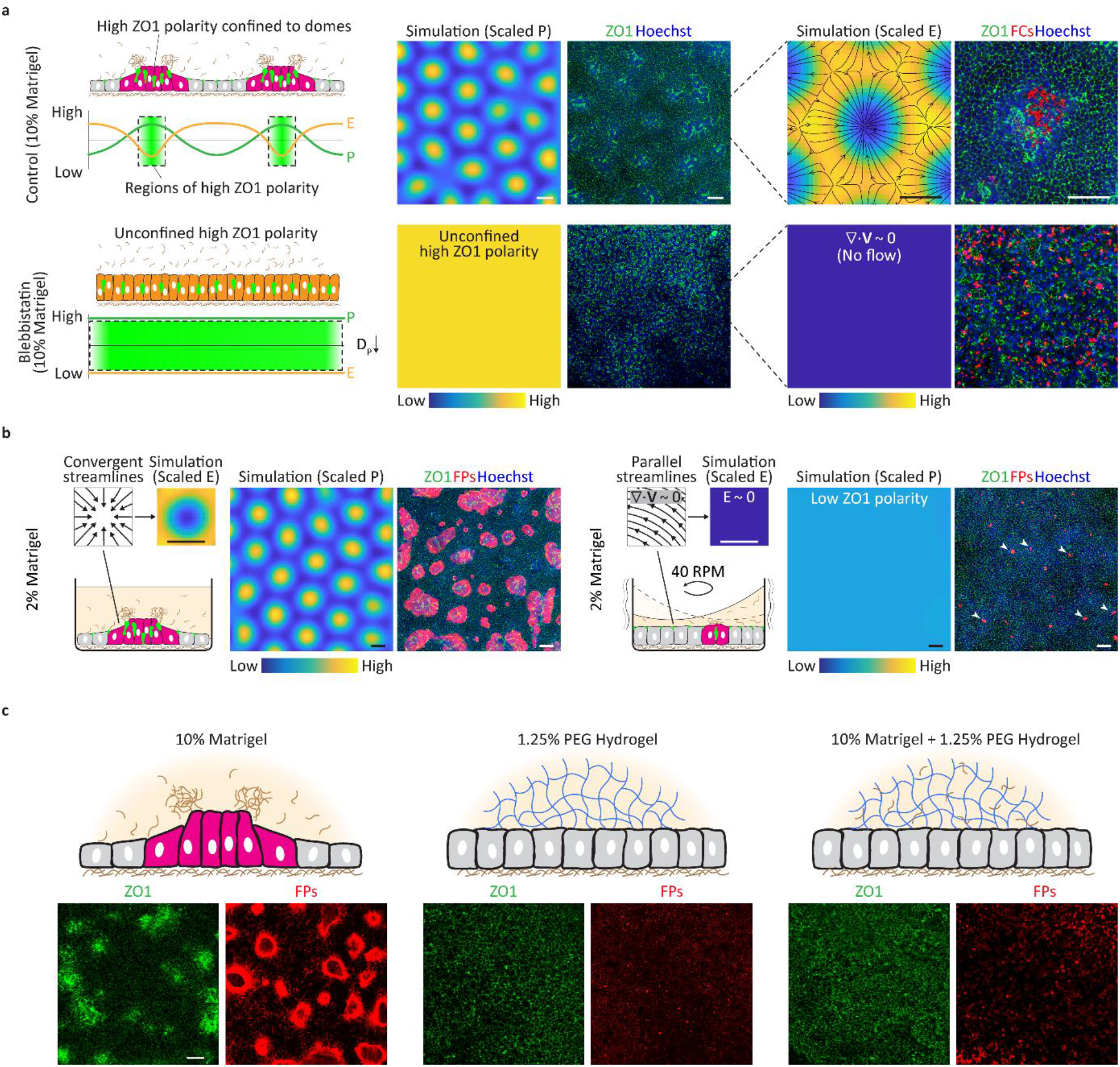
Polarity, ECM flow and ECM mechanics perturbations and effects on morphogenesis and flow establishment. **a** Schematic representation of dome morphogenesis and ECM flow relation in intact and impaired polarity and accompanied respective scale E and P profiles. Representative scaled P with corresponding ZO1 expression at 24 h in intact and impaired (10 μM Blebbistatin) polarity conditions (n = 3). Scaled E with corresponding ZO1 expression and FC distribution (n = 3). Scalebars 100 μm. **b** Schematic representation of dome morphogenesis and ECM flow under stable conditions, with accompanied representative streamlines and related scaled E values (n = 3). Scaled P values with corresponding ZO1 and FP distribution at 24 h in 2% Matrigel conditions. Schematic representation of dome morphogenesis and ECM flow under shaking (40 RPM) conditions, with accompanied representative streamlines and related scaled E values (n = 3). Scaled P values with corresponding ZO1 and FP distribution at 24 h in 2% Matrigel under shaking (40 RPM) conditions (n = 3). Scalebars 100 μm. **c** Schematic representation of Matrigel condition. Representative image of ZO1 expression and FP distribution at 24 h in 10% Matrigel conditions (n = 3). Schematic representation of PEG (blue lines) condition. Representative image of ZO1 expression and FP distribution at 24 h in 10% Matrigel conditions (n = 3). **c** Schematic representation of PEG-Matrigel condition. Representative image of ZO1 expression and FP distribution at 24 h in 10% Matrigel conditions (n = 3). Scalebars 100 μm.

**Figure 4.**
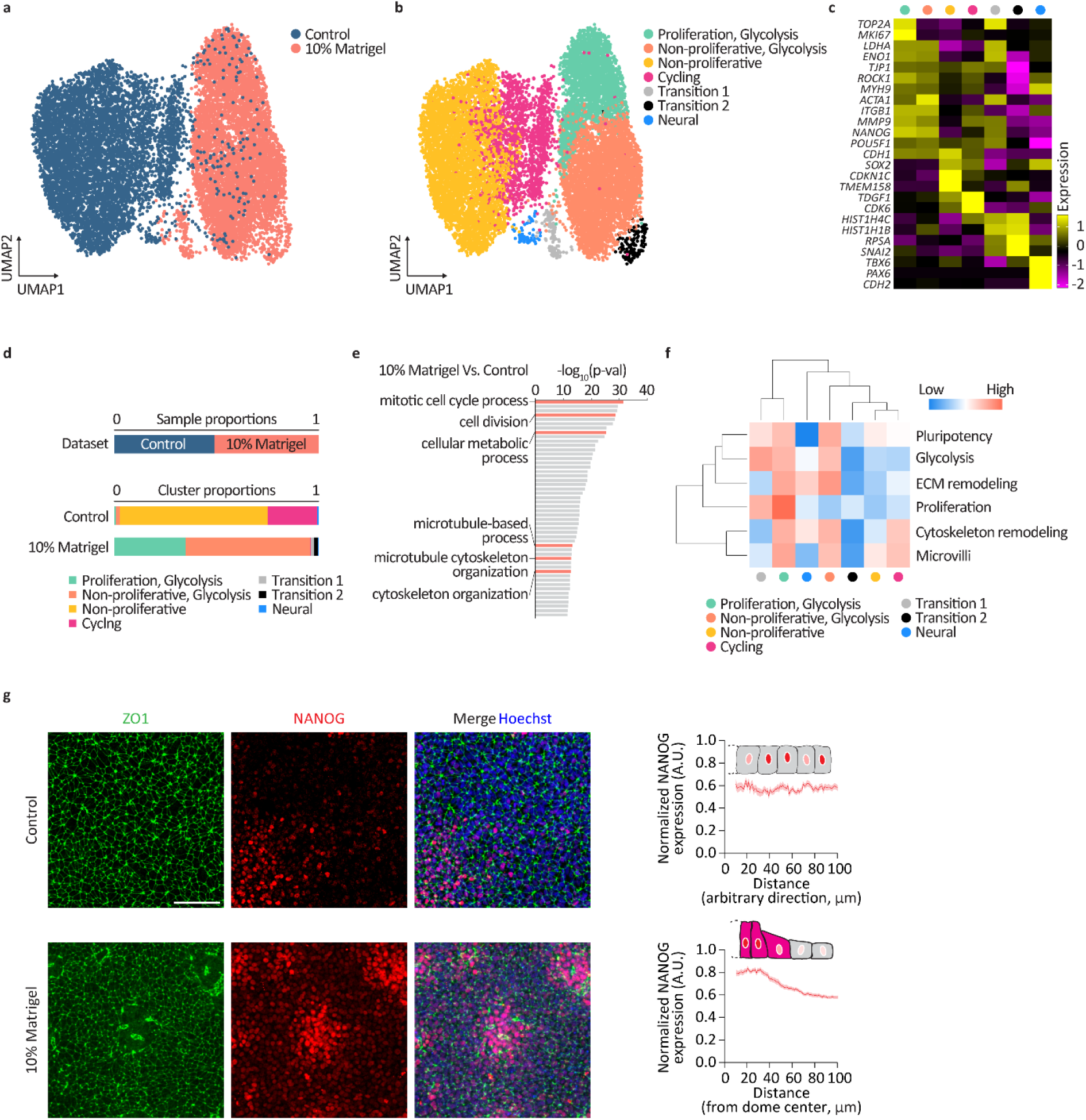
Single cell RNA-seq reveals the transcriptomic landscape of hPSCs in control and 10% Matrigel conditions. **a** UMAP of the combined dataset (control and 10% Matrigel). **b** UMAP of the combined dataset with identified clusters (Proliferative cells with glycolysis activity (Proliferation, Glycolysis), cell with glycolysis activity (Glycolysis), non-proliferative cell (Non-proliferative), cycling cells (Cycling), transitioning cells 1 (Transition 1), transitioning cell 2 (Transition 2), neural differentiating cells (Neural). Top marker genes for each cluster in Supplementary Data 1. **c** Exclusive expression in specific clusters of selected marker genes for proliferation, cytoskeletal, extracellular matrix remodeling, pluripotency, cell cycling and senescence, early mesoderm and neuroectoderm identities. Expression values are normalized and centered. **d** Dataset sample proportion and sample cluster proportions. **e** GO enrichment analysis of upregulated key processes in 10% Matrigel conditions compared to control conditions include proliferation, metabolic activity, and cytoskeletal and microtubule organization. **f** Hierarchical clustering of key processes and cellular states based on their gene-set scores for each cluster. Expression values are row-scaled. **g** Representative images ZO1 and NANOG expressions in control and 10% Matrigel conditions at 24 h (n = 3) with corresponding normalized NANOG radial profiles (n = 3 for a total of 30 domes). Scalebars 100 μm.

The model equally predicted the importance of a stable ECM flow with divergent characteristics to support high polarity regions (**Fig.4b**). To investigate this further, we aimed to perturb naturally occurring flow divergence patterns by introducing exogenous flows to the culture through an orbital shaker. Upon shaking, a global flow was imposed (parallel streamlines and divergence, ∇∙**V** = 0), resulting in a single region of low ZO1 polarity. Here, instead of Matrigel toroid formation, weak accumulation of ECM was still possible, likely due to the no-slip fluid boundary condition at the Matrigel-monolayer interface, (**Fig.4b**, white arrowheads). This helped stabilize scattered symmetry breaking events (**Fig.4b**, white arrowheads), but was insufficient to mature them into domes. Therefore, unperturbed ECM-flow is required for dome morphogenesis and relies on transport of the matrix.

To further investigate the requirement of ECM flow on dome morphogenesis we replaced the matrix with a polyethylene glycol (PEG) hydrogel matrix, which is known to support morphogenesis^9^ and can deform, but which is linear elastic, nondegradable, and cannot flow. In standard Matrigel conditions, flow was observed at the cell-Matrigel interface accompanied by localized high expressions of ZO1 (**Fig.4c**). However, in PEG and PEG supplemented with Matrigel conditions, neither ECM flow nor dome morphogenesis were observed, as evidenced by homogenous ZO1 expression and unaltered FP distributions (**Fig.4c**), indicating that Matrigel components alone are insufficient to promote morphogenetic events and that preventing the flow capability of the ECM results in morphogenetic failure.

Overall, *in silico* simulations and complementary *in vitro* observations argue for a strongly coupled feedback between ECM flow and morphogenesis: while ECM flow is triggered by symmetry-breaking events, the maintenance and maturation of such events relies on the establishment of the flow.

## The transcriptomic landscape of hPSCs in Matrigel and control conditions

To unravel transcriptomic changes in hPSCs upon exposure to Matrigel, we performed scRNAseq on hPSCs in control and 10% Matrigel conditions after 24 h of Matrigel exposure. Graph-based clustering of the 13,250 cells retained for analysis displayed a stark contrast between control cells and those exposed to 10% Matrigel for 24 h (**Fig.5a**). Based on upregulated genes as well as specific markers, seven clusters were identified and annotated to reveal proliferative cells with glycolysis activity (Proliferation, Glycolysis), non-proliferative cells with glycolysis activity (Non-proliferative, Glycolysis), non-proliferative cells with oxidative activity (Non-proliferative), cycling cells (Cycling), neural differentiating cells (Neural), and two transition clusters with various marker genes (Transition 1/2) (**Fig.5b,c**). The dataset was equally distributed between the control (49%) and Matrigel (51%) conditions (**Fig.5d**). However, the control condition largely comprised cycling and non-proliferative cells, whereas exposure to Matrigel prompted proliferation, and glycolysis activity. This finding was supported by the gene ontology (GO) enrichment analysis where cell proliferation and metabolic processes were upregulated in Matrigel condition (**Fig.5e**), in addition to those involved in microtubule and cytoskeletal organization. In addition to glycolysis and proliferation genesets, pluripotency was upregulated in clusters belonging to the Matrigel condition (**Fig.5f**). This was confirmed by NANOG staining, most abundantly expressed in the domes of the Matrigel conditions, (**Fig.5g**).

## Microvilli drive ECM Flow

Since directed ECM flow is driven by the apical surface of epithelial monolayers, we investigated whether surface microstructures such as microvilli would be involved in ECM transport. The role of microvilli was particularly promising due to their link with cytoskeleton remodeling^15,16^, a process which is upregulated in Matrigel conditions. Indeed, we found the microvilli and cytoskeletal remodeling genesets cluster together, with the former expressed higher in Matrigel clusters compared to control ones (**Fig.5f**). These results suggest that cytoskeletal remodeling and associated microvilli dynamics may drive ECM flow. It is known that microvilli biogenesis and function is triggered by shear forces from flows^17^. Moreover, microvilli have been shown to exhibit motion^18^ and active mobility across apical membranes^19^ of epithelial cells. We therefore used scanning electron microscopy (SEM) to reveal changes in surface and microvilli morphology upon Matrigel exposure. Microvilli were sparsely populated and had arbitrary orientation on the apical surfaces of cells in control conditions (**Fig.6a**), while in cells exposed to Matrigel, microvilli appeared densely populated and were clearly excluded from dome regions where Matrigel could be seen (**Fig.6a**, red arrow heads). Interestingly, Matrigel bundles projected radially out of dome regions while surrounding microvilli were coherently oriented towards the domes (**Fig.6a**), suggesting that collective and directional microvilli motion is used to transport ECM.

**Figure 6.**
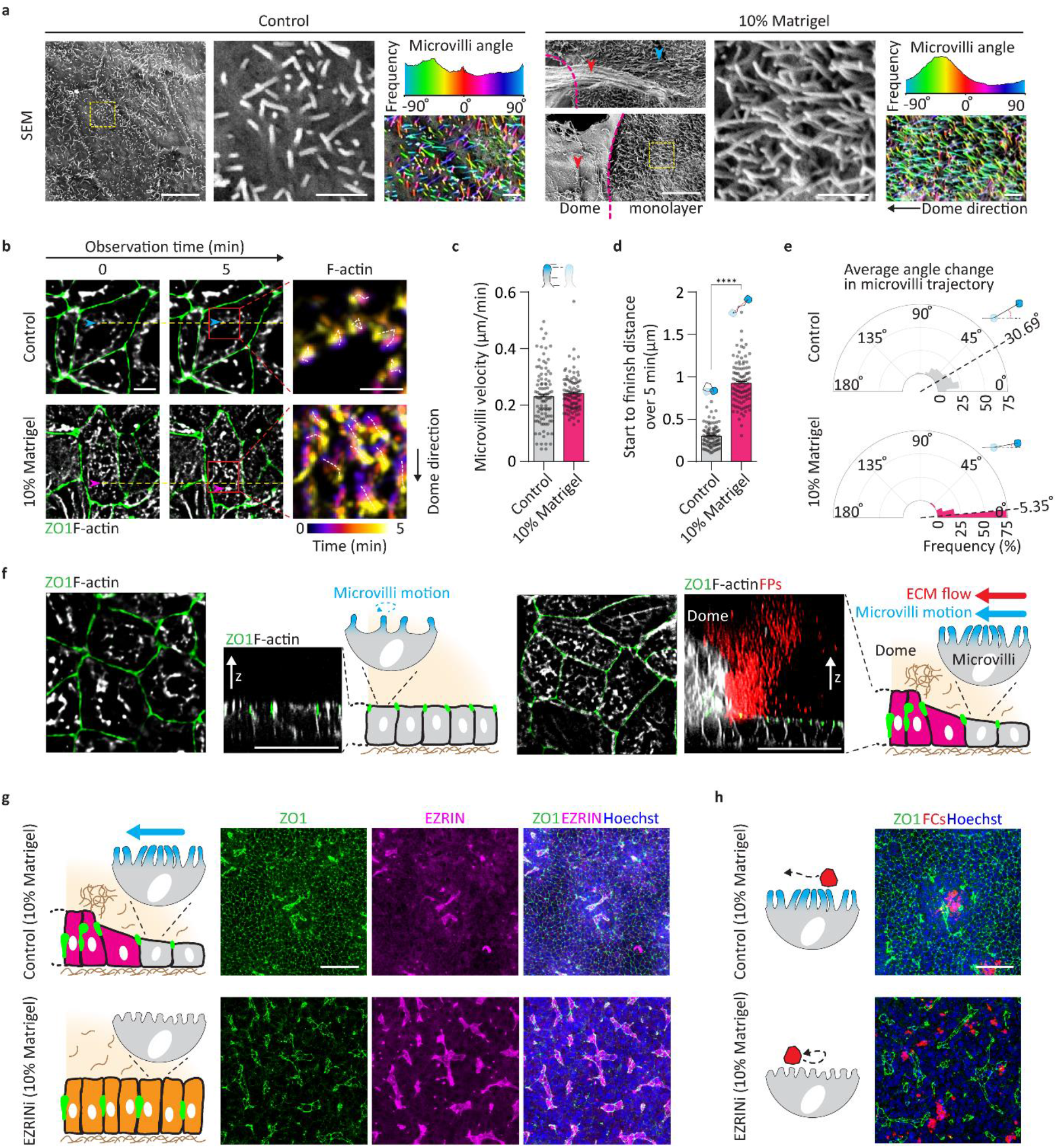
Microvilli dynamics drive ECM flow. **a** Scanning electron microscopy images at 24 h of cells under control and 10% Matrigel images (n = 1 with technical triplicates). Scalebars 5 μm (insets scalebars 2 μm). Microvilli orientation angel frequency color coded for angle (−90 – 90°) with accompanied representative microvilli image with superimposed orientation angle color. (n = 1 with technical triplicates). Scalebars 2 μm. **b** Live F-actin and ZO1 dynamics in control and 10% Matrigel conditions at 24 h. Surface F-actin dynamics color coded for time (n = 3). **c** Measured microvilli (F-actin) velocities over 5 min in control and 10% Matrigel conditions (n = 3 for a total of 100 F-actin foci). Error bars are SEM. **d** Measured microvilli (F-actin) trajectory start to finish distances over 5 min in control and 10% Matrigel conditions (from **c**). Statistical analysis was determined by unpaired two-sided t-test *p* < 0.0001. Error bars are SEM. **e** Measured microvilli (F-actin) trajectory average angle change over 5 min in control and 10% Matrigel conditions (from **c**). **f** Top and side view representative images of ZO1 and F-actin expressions in control and 10% Matrigel conditions with FP distribution at 24 h. Schematic representation of microvilli dynamics in control and Matrigel conditions. **g** Schematic representation of microvilli dynamics in control 10% Matrigel and under EZRIN inhibition (5 μM EZRINi - NSC668394) 10% Matrigel conditions and corresponding representative images of ZO1 and EZRIN expressions (n = 3). Scalebars 100 μm. **h** Corresponding FCs localization in 10% Matrigel control and EZRINi conditions. Scalebars 100 μm.

To reveal microvilli motion, we investigated F-actin dynamics at the apical side of monolayer cells surrounding domes, as F-actin is known to anchor to microvilli^18,19^ (**Fig.6b**, **Supplementary Movie 5**). Compared to control conditions, exposure to Matrigel resulted in similar microvilli velocities (**Fig.6c**) but with clearly linear and non-stochastic trajectories (**Fig.6a**), longer start-to-finish distances (**Fig.6d**) and smaller changes in trajectory angles (**Fig.6e**). Overall, measured microvilli velocities closely matched previous studies ~0.200 μm/min^19^, and corresponded well with measured ECM flow velocities of ~0.200 μm/min of zone 1 at 24 h (**Fig.1d**). The results suggest that upon exposure to ECM components, the sparse population of microvilli undergo increased biogenesis and acquire coherent translational motion directed towards domes, which likely drags and help accumulate ECM around dome positions.

To further demonstrate the role of microvilli microstructures on dome morphogenesis and ECM flow generation, we impaired EZRIN, a linker protein between F-actin and the microvillus structure. Compared to control conditions, exposure to EZRIN inhibitor (EZRINi, NSC668394) abrogated dome morphogenesis as well as control-like monolayers surrounding the domes (**Fig.6g**). In this case EZRIN expression became confined only to high ZO1 polarity regions and was not extended to the surrounding tissue as in EZRINi-free conditions (**Fig.6g**). With impaired functionality of microvilli upon EZRINi treatment, cell monolayers were unable to drive ECM flow leaving scattered FCs across the cellular surface compared to EZRINi-free conditions where clear FC aggregation is observed at dome locations (**Fig.6h**).

Overall, these results suggest that coherent microvilli dynamics are involved in guiding ECM flow towards sites of morphogenesis. In this way, microvilli directly encode morphogenetic events into mechanical flows, which in turn define multicellular morphogenesis across the tissue.

## ECM flow-morphogenesis interplay defines the patterning landscape

In addition to morphogenetic processes, we also explored whether ECM flow could mediate the specification of cell fates. To gain insight into the cell fates generated upon ECM exposure in our system, we mapped our scRNAseq dataset to that of a human gastrula (E-MTAB-9388). By examining the top three ranked predictions, we found, as expected, that both conditions resembled most the epiblast (EP), the undifferentiated and earliest form of a gastrulating embryo (**Fig.7a,b**). However, unlike in control condition, exposure to ECM flow gave rise to more primitive streak (PS)-like and nascent mesoderm (NM)-like cells than in control conditions (**Fig.7a,b, c**). These results suggest that exposure to Matrigel and ECM flow promoted a transition from the EP state into the PS and NM. We reasoned that this bias may facilitate mesoderm differentiation and pattering and therefore focused on the expression of brachyury (T), which is associated with epiblast^20^ and primitive streak^21^ formation as well as mesoderm differentiation^22^ and somitogenesis^12^.

**Figure 7.**
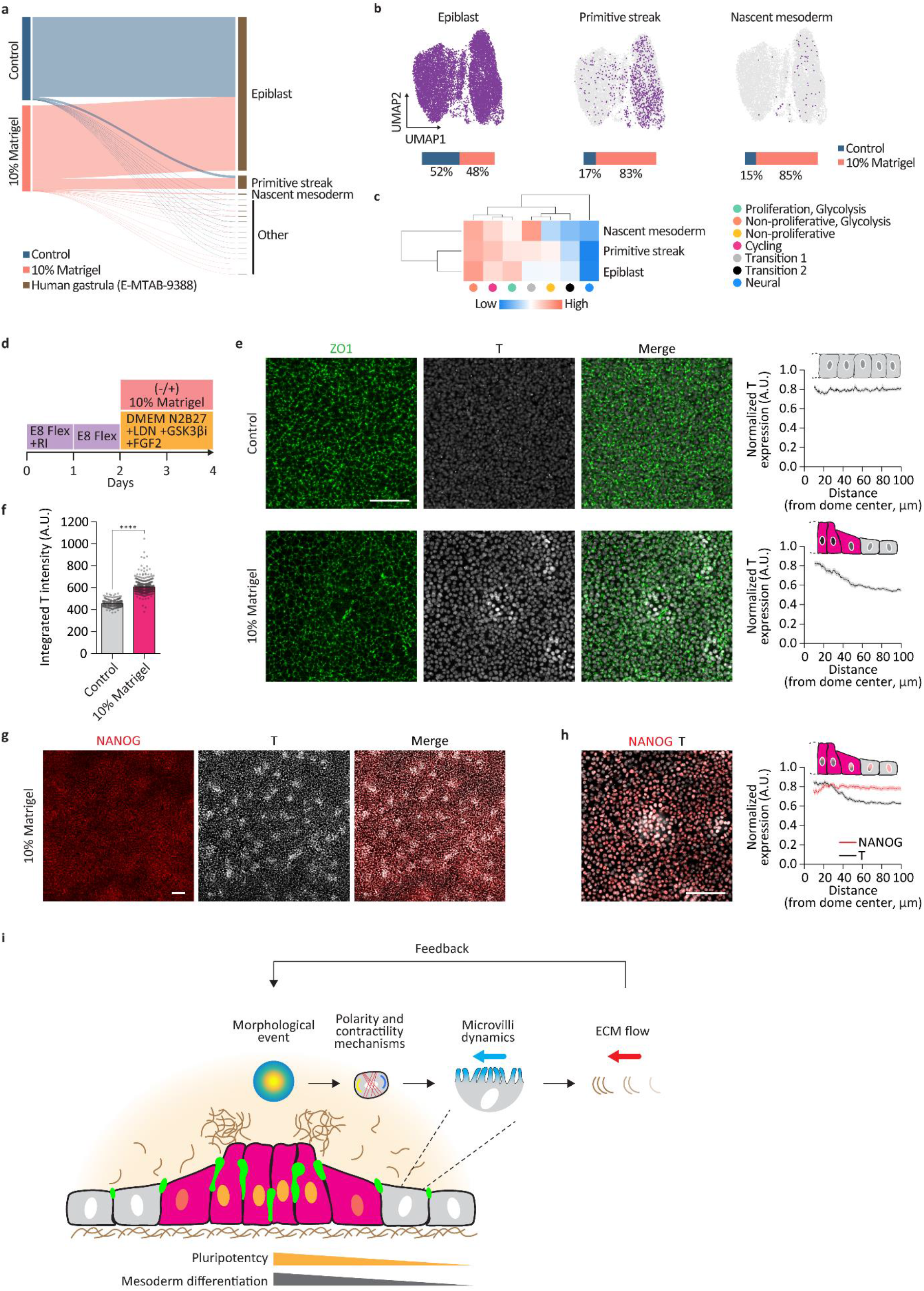
ECM flow and morphogenesis relation defines the cell patterning landscape. **a** Mapping of dataset to the human gastrula dataset (E-MTAB-9388). **b** UMAPs revealing cells categorized as predicted epiblast, primitive streak, and nascent mesoderm after dataset mapping. Accompanied sample proportion for each prediction. **c** Hierarchical clustering of epiblast, primitive streak, and nascent mesoderm based on their gene-set scores for each cluster. Expression values are row-scaled. **d** Mesoderm differentiation protocol. **e** Representative images of endpoint day 4 expressions of ZO1 and T (n = 3) in control and 10% Matrigel conditions with corresponding ZO1 radial profiles (n = 3 for a total of 30 profiles per condition). **f** Integrated intensity of T expression in individual cells using consecutive confocal images (z-stack) in control and 10% Matrigel conditions (n = 3 for a total of 450 cells per condition). **g** Representative images of endpoint day 4 expressions of NANOG and T in 10% Matrigel condition (n = 3). **c** Representative images of NANOG and T in 10% Matrigel condition (n = 3) with corresponding NANOG and ZO1 radial profiles (n = 3 for a total of 30 domes). **d** Proposed model for ECM flow-morphogenesis reciprocal interplay. Scalebars 100 μm.

To differentiate epithelial monolayers, we cultured hPSC in a 48 h mesoderm differentiation protocol, using a combination of BMP and WNT pathway modulation and exposure to bFGF and 10% Matrigel (**see methods**, **Fig.7d**). Although highly defined dome structures were less obvious than under pluripotent conditions, they retained ZO1 polarity and were associated with high T expression, compared to homogeneous expressions in control conditions (**Fig.7e**). Overall, T expression intensity was higher upon Matrigel exposure (**Fig.7f**), suggesting that the presence of ECM enhances mesoderm differentiation, in accordance with transcriptomic analysis (**Fig.7a-c**). The patterned T expressions in Matrigel conditions contrasted with the relatively homogeneous NANOG fate (**Fig.7g,h**), suggesting that feedback mechanisms between ECM flows and morphogenesis help define the early patterning landscape during human gastrulation.

Overall, we show that morphogenetic symmetry breaking events encode the conversion of polarity signals to mechanical ECM flows (**Fig.7i**), and that this process is mediated by the directed motion of microvilli. In doing so, a strong feedback between morphogenesis and ECM flow is generated, which if interrupted, leads to failure in morphogenesis. Through an *in silico* model, we show this ECM-tissue interplay follows reaction-diffusion-like principles, rendering this a periodic and tissue-wide phenomenon, where cell polarity anticorrelates with flow divergence. In this way we recapitulate and predict observed flow and morphogenesis patterns, which may be extendable to previously reported ECM-driven and micropattern-mediated morphogenetic events, such as neural tube morphogenesis^13^. Indeed, such cases may be specific, geometrically constrained solutions to the generalized, geometrically unconstrained ECM flow-driven phenomena we describe here. It remains to be investigated whether actin treadmilling drives ECM flow and whether the coordination and coherence in ECM dynamics is guided by tissue tension gradients. The observed expressions of NANOG and T during dome formation models early development processes at the onset of, and during, gastrulation. While bulk ECM motion^6^ and ECM-tissue coupling^23^ have been reported *in-vivo*, our *in-vitro and in-silico* results reveal a previously unrecognized role for ECM transport and flow feedback to support morphogenesis and cell patterning. These results establish ECM flow as a novel guiding mechanism during development and as a driver of epithelial morphogenesis. With the increasing use of ECMs in tissue and organoid culture^10–13^ our approach can be expanded to help unravel the role of ECM flow in guiding morphogenesis in other model systems, in health and disease settings.

## Supplementary figures

**Supplementary Figure 1.**
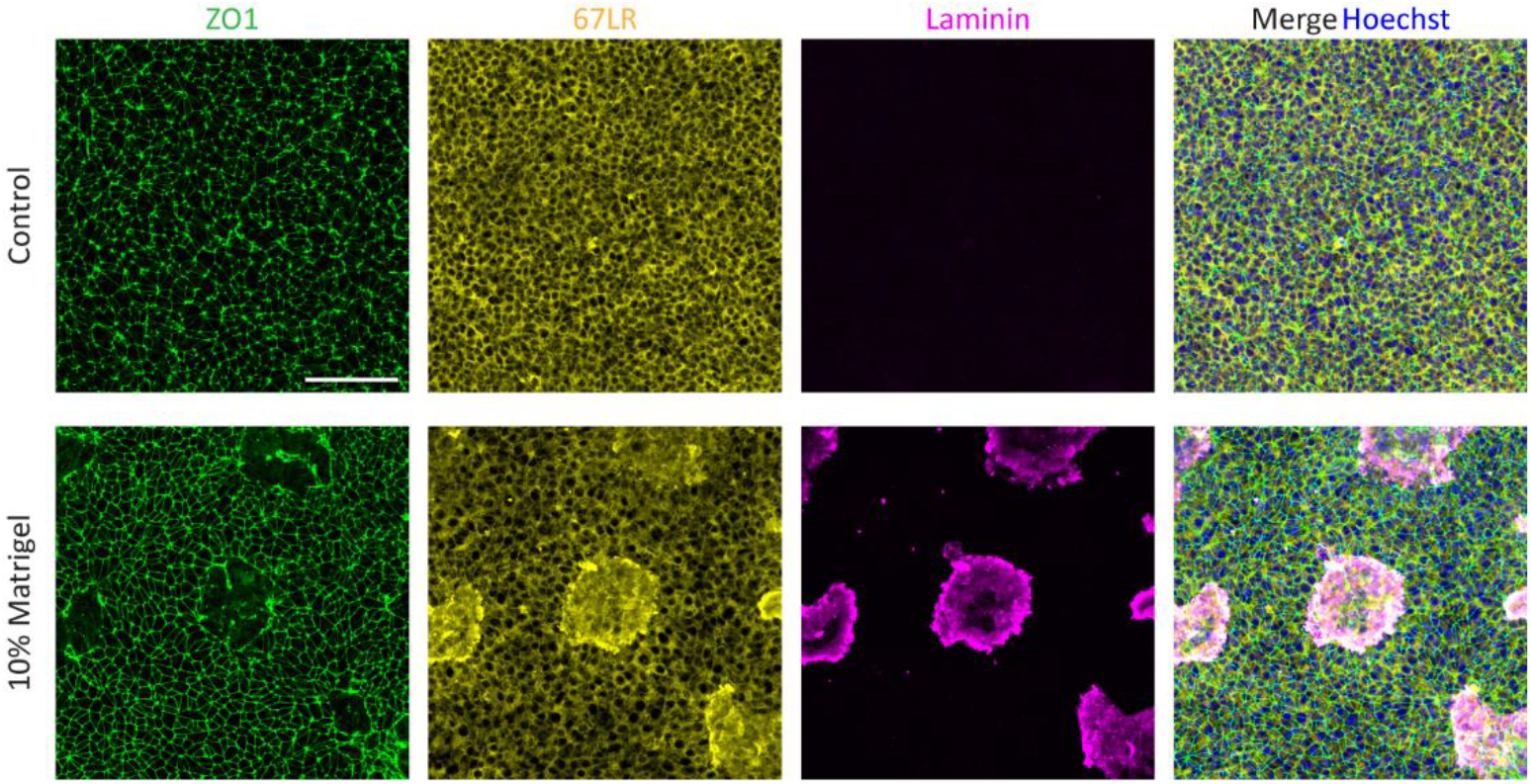
Matrigel localization affects laminin receptor expression. Representative images of ZO1, 67LR and laminin expressions in control and 10% Matrigel conditions at 24 h. Scalebars 100 μm.

**Supplementary Figure 2.**
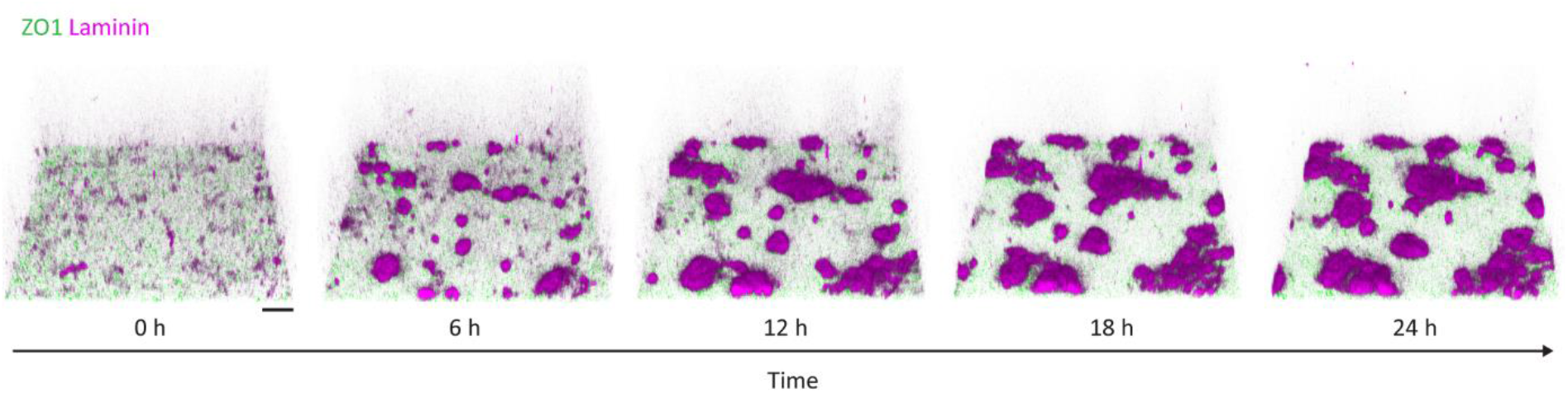
Laminin dynamics over 24 h. Representative 3-D reconstructed images of ZO1 and laminin temporal dynamics over 24 h (n = 3). Scalebars 100 μm.

**Supplementary Figure 3.**
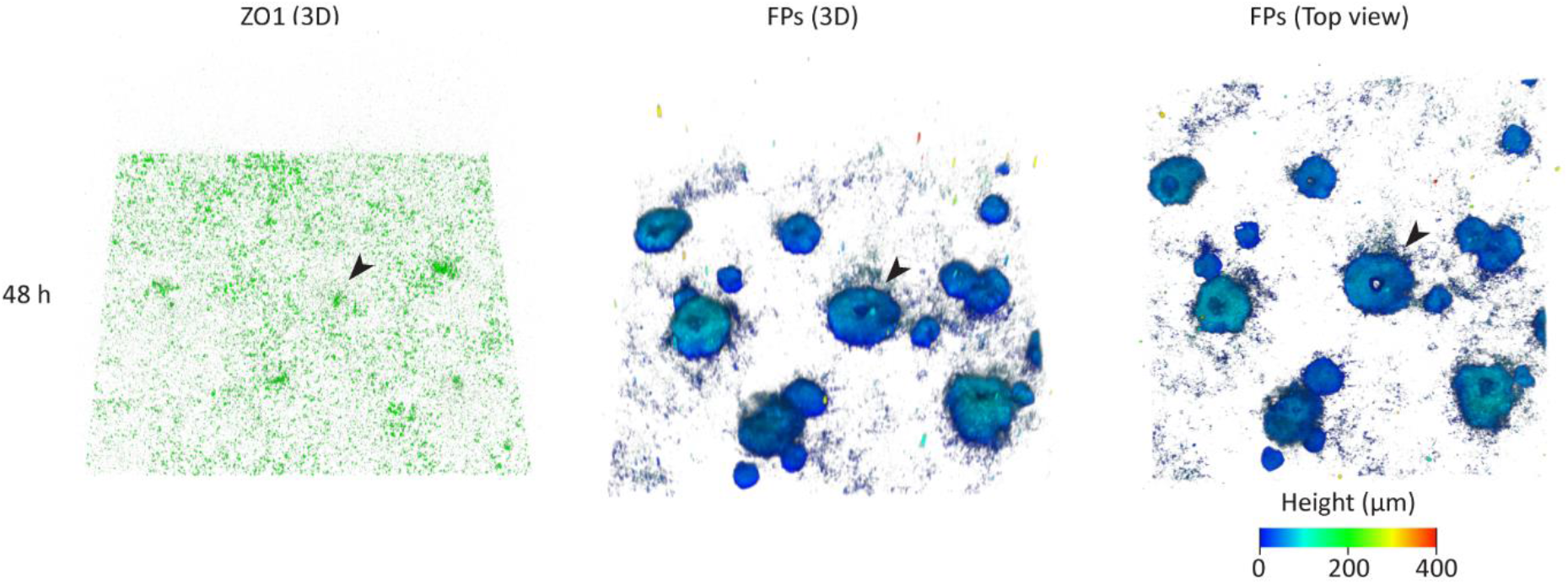
Matrigel toroid formation at 48 h. Representative 3-D reconstructed image of ZO1 expression and Matrigel-tagged FPs at 48 h. Top view of 3-D reconstructed image of Matrigel-tagged FPs (n = 2). FPs channel color coded for height. Black arrow point to a representative dome formation event. Scalebar 100 μm.

**Supplementary Figure 4.**
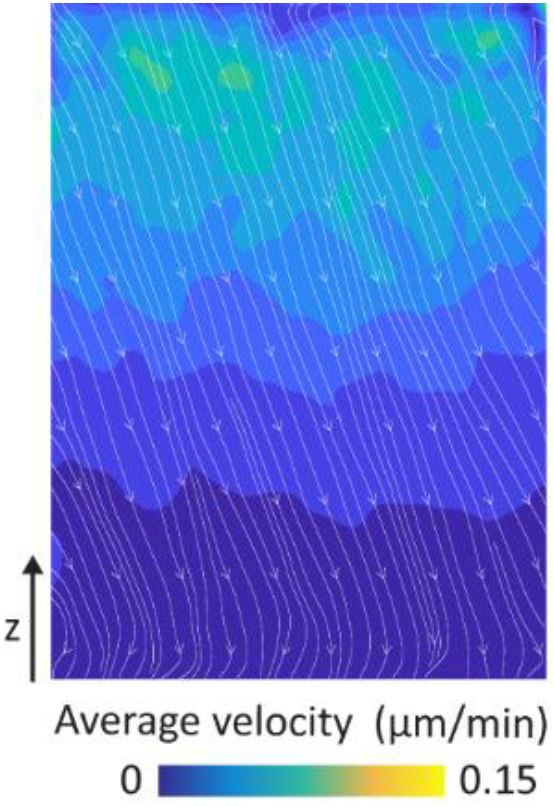
ECM dynamics in the absence of cells. 24 h average velocity map color coded for velocity magnitude where white lines and arrowhead denote flow streamlines and direction. The recorded velocities in control conditions can be attributed to the downward ECM flow direction due to settling during Matrigel polymerization. (n = 4).

**Supplementary Figure 5.**
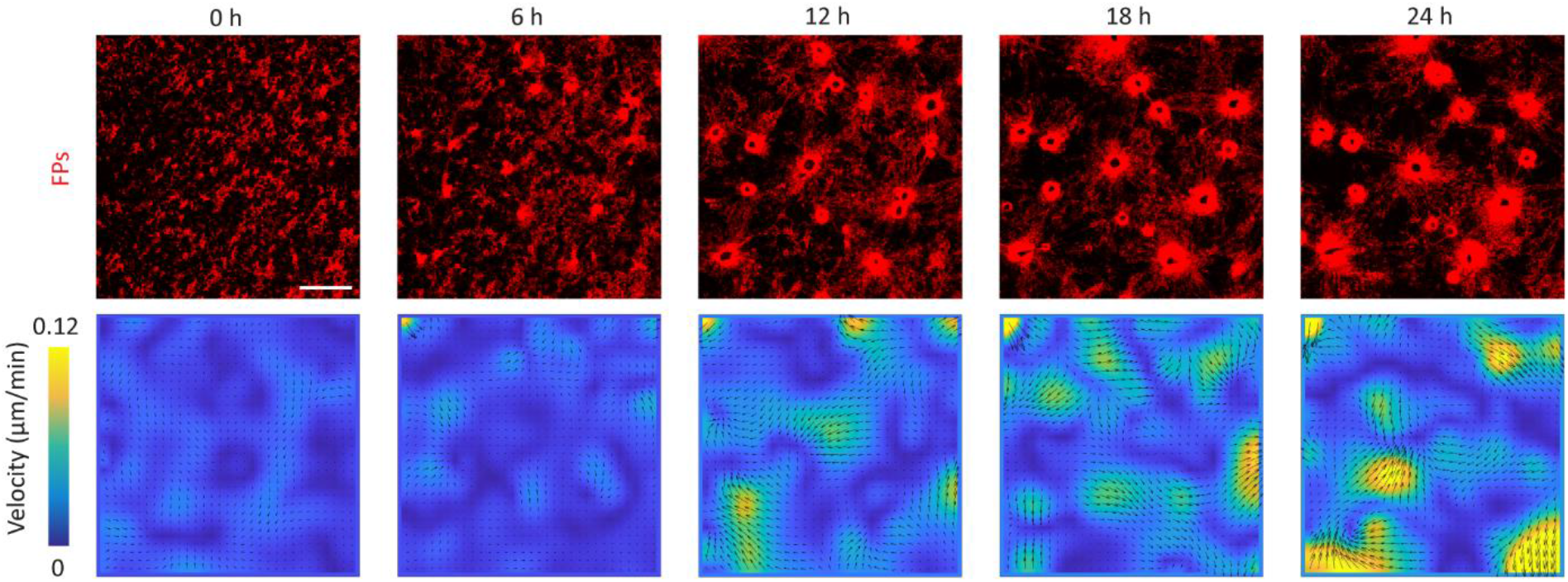
FP distribution and corresponding ECM flow field. FP distribution over 24 h with corresponding instantaneous velocity map color coded for velocity magnitude where black arrows depict velocity vector.

**Supplementary Figure 6.**
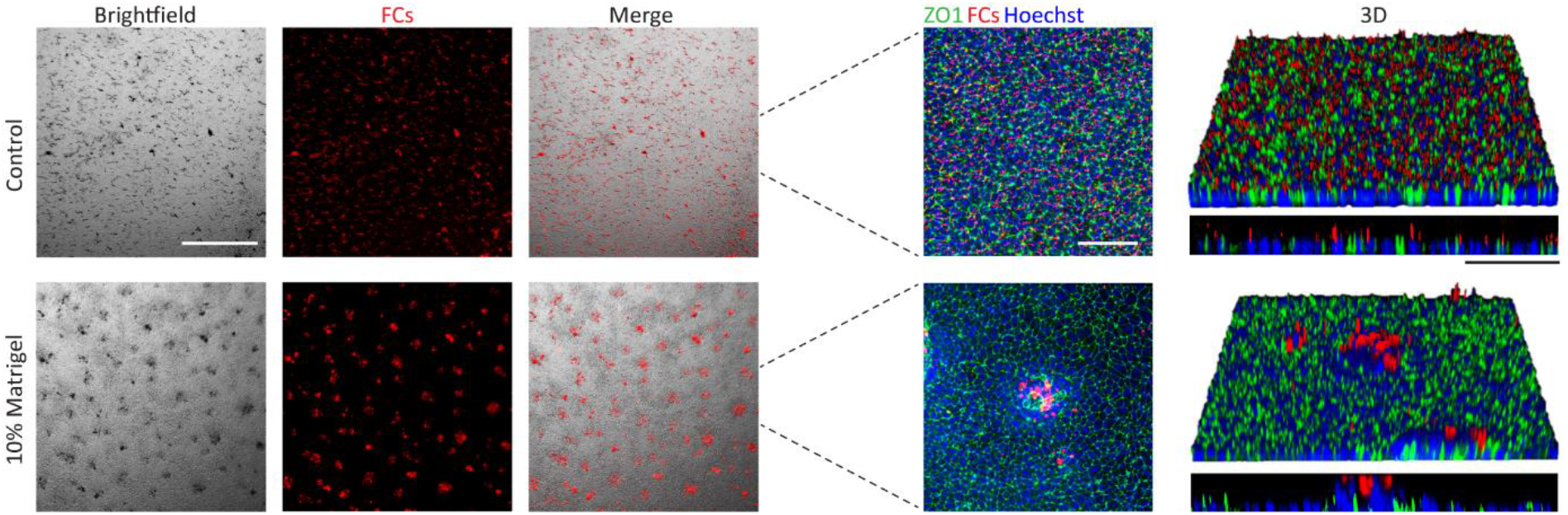
FC localization in 10% Matrigel conditions. Representative low magnification images of control and 10% Matrigel conditions showing FCs distribution (n = 3). Scalebar 500 μm. Representative images of control and 10% Matrigel conditions with accompanied 3D reconstructed and side view images (n = 3). Scalebar 100 μm.

**Supplementary Figure 7.**
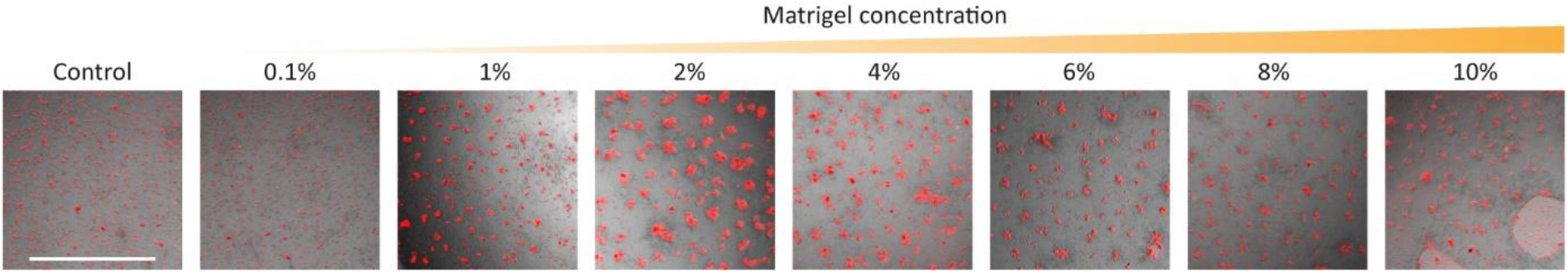
FC localization across different Matrigel conditions. Representative low magnification images FCs distribution across various concentrations (n = 3). Scalebar 500 μm.

**Supplementary Figure 8.**
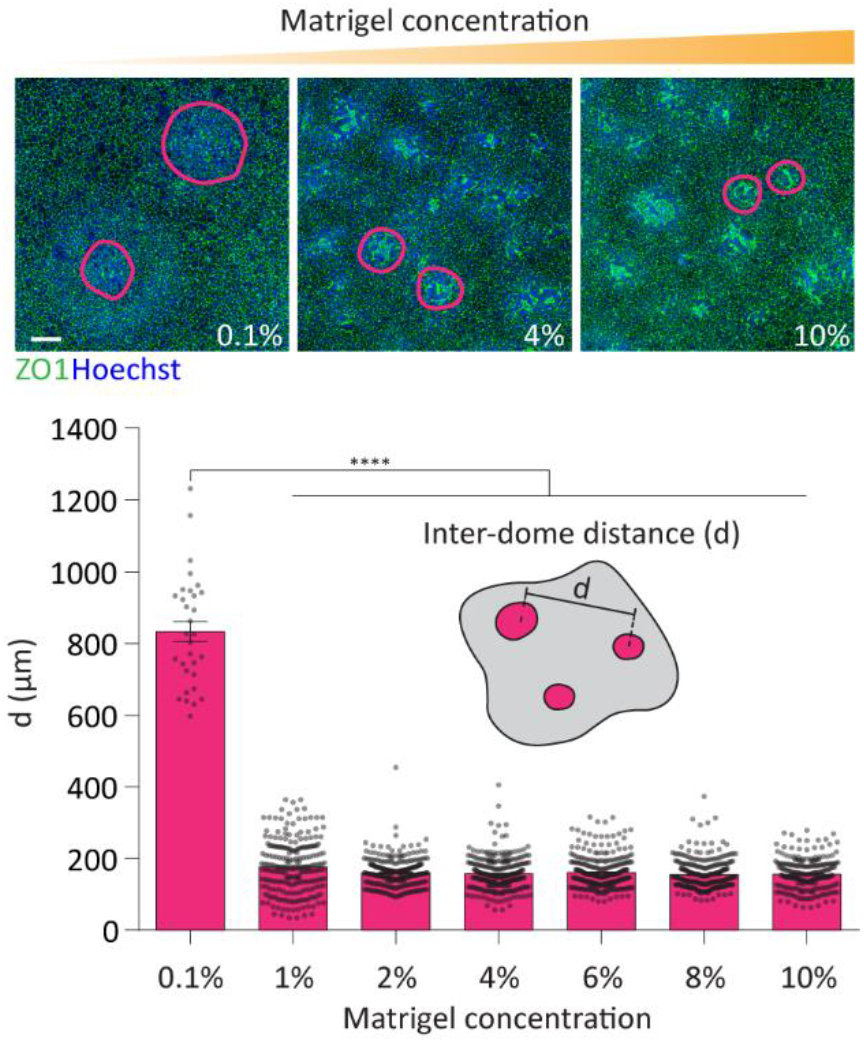
Matrigel concentration modulates dome periodicity. Representative images of various Matrigel conditions at 24 h showing nuclei and ZO1 expression (n = 5), example dome locations are encircled. Quantification of inter-dome distance at various Matrigel conditions (n = 5 for >200 domes per conditions except for 0.1% Matrigel (31 domes), statistical analysis was determined by one-way ANOVA for multiple comparisons where the 0.1% Matrigel case had *p*-value < 0.0001 compare to all other conditions.

**Supplementary Figure 9.**
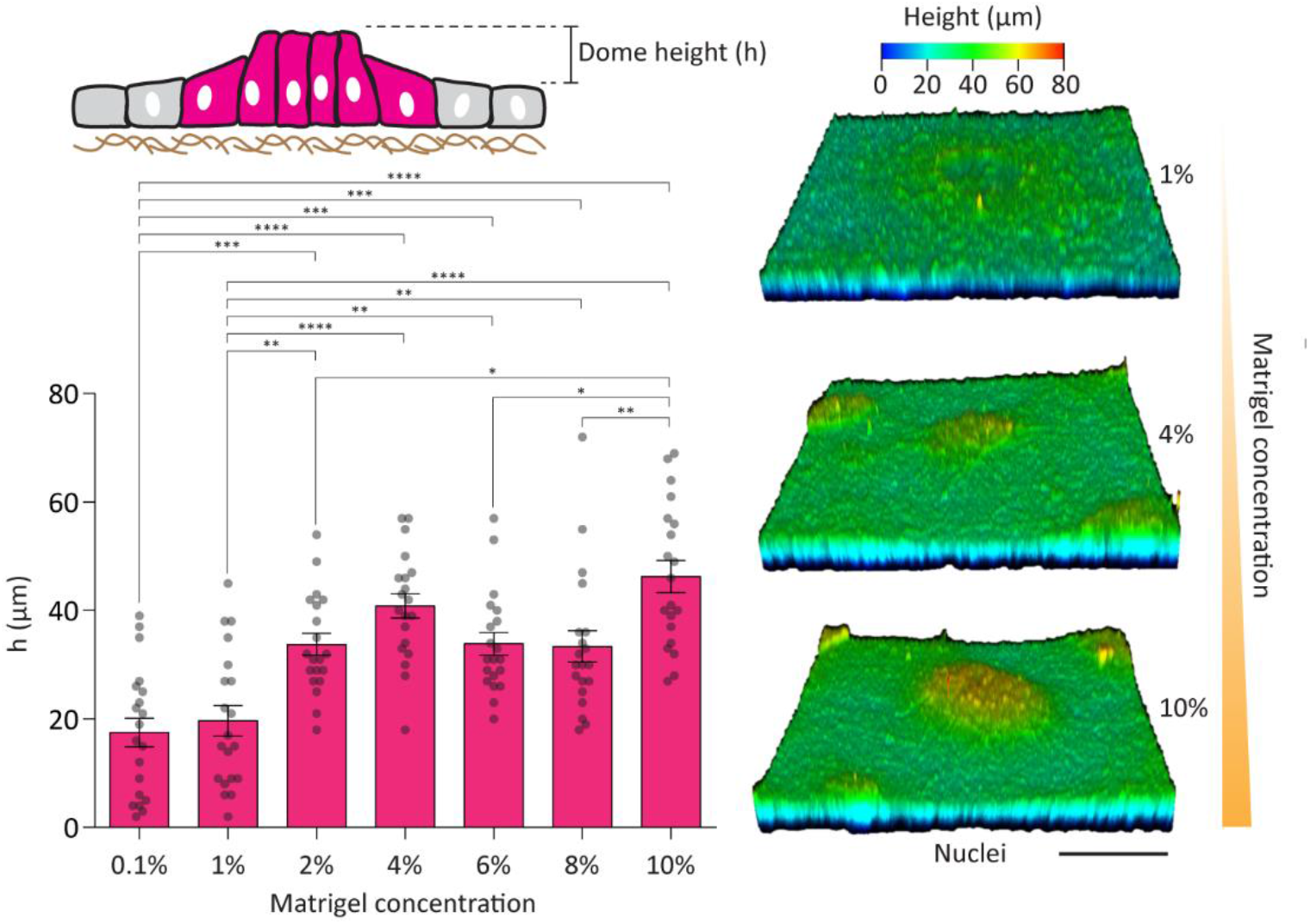
Matrigel concentration modulates dome height. Schematic representation of dome morphogenesis and height. Quantification of dome height for various Matrigel conditions (n = 3 for a total of 20 domes per condition). Statistical analysis was determined by one-way ANOVA for multiple comparisons 0.1-2% *p* = 0.0003, 0.1-4% *p* < 0.0001, 0.1-6% *p* = 0.0003, 0.1-8% *p* = 0.0005, 0.1-10% *p* < 0.0001, 1-2% *p* = 0.0029, 1-4% *p* < 0.0001, 1-6% *p* = 0.0026, 1-8% *p* = 0.0042, 1-10% *p* < 0.0001, 2-10% *p* = 0.0128, 6-10% *p* = 0.0140, 8-10% *p* = 0.0089. Representative 3-D reconstructed images showing nuclei position color coded for height (n = 3).

**Supplementary Figure 10.**
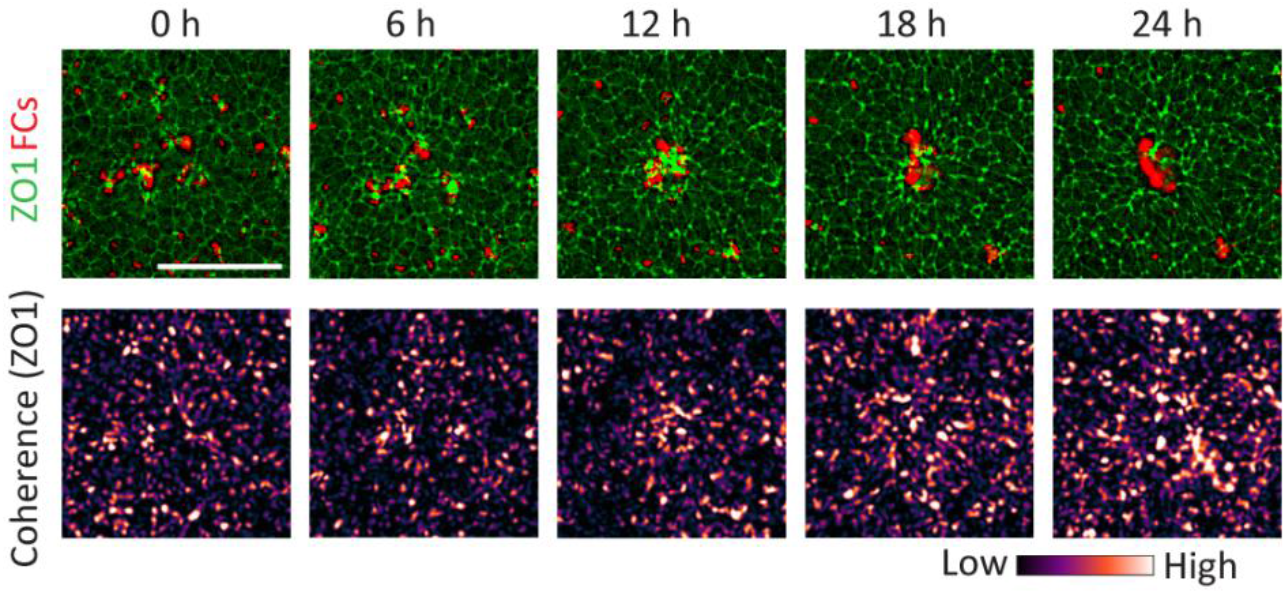
Coherence dynamics in 10% Matrigel conditions. Representative images showing ZO1 expression, FC localization and Coherence score dynamics over 24 h. Scalebar 100 μm.

**Supplementary Figure 11.**
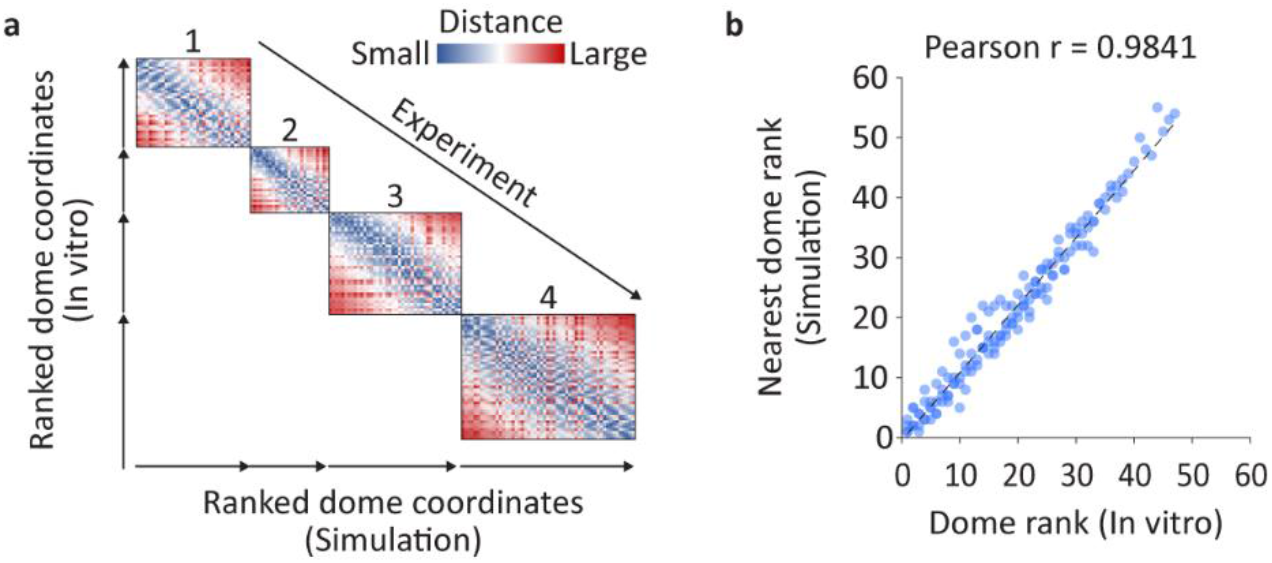
Correlation between *in silico* and *in vitro* dome positions. **a** Heatmap showing ranked *in vitro* and *in silico* domes and the distance between them in a shared space color coded for the separation length (from **Fig.3e**). **b** Correlation between ranked *in vitro* domes and respective nearest *in silico* domes, *p*-value denotes Pearson correlation analysis *p* < 0.0001 (n = 4 for 143 dome pairs).

**Supplementary Figure 12.**
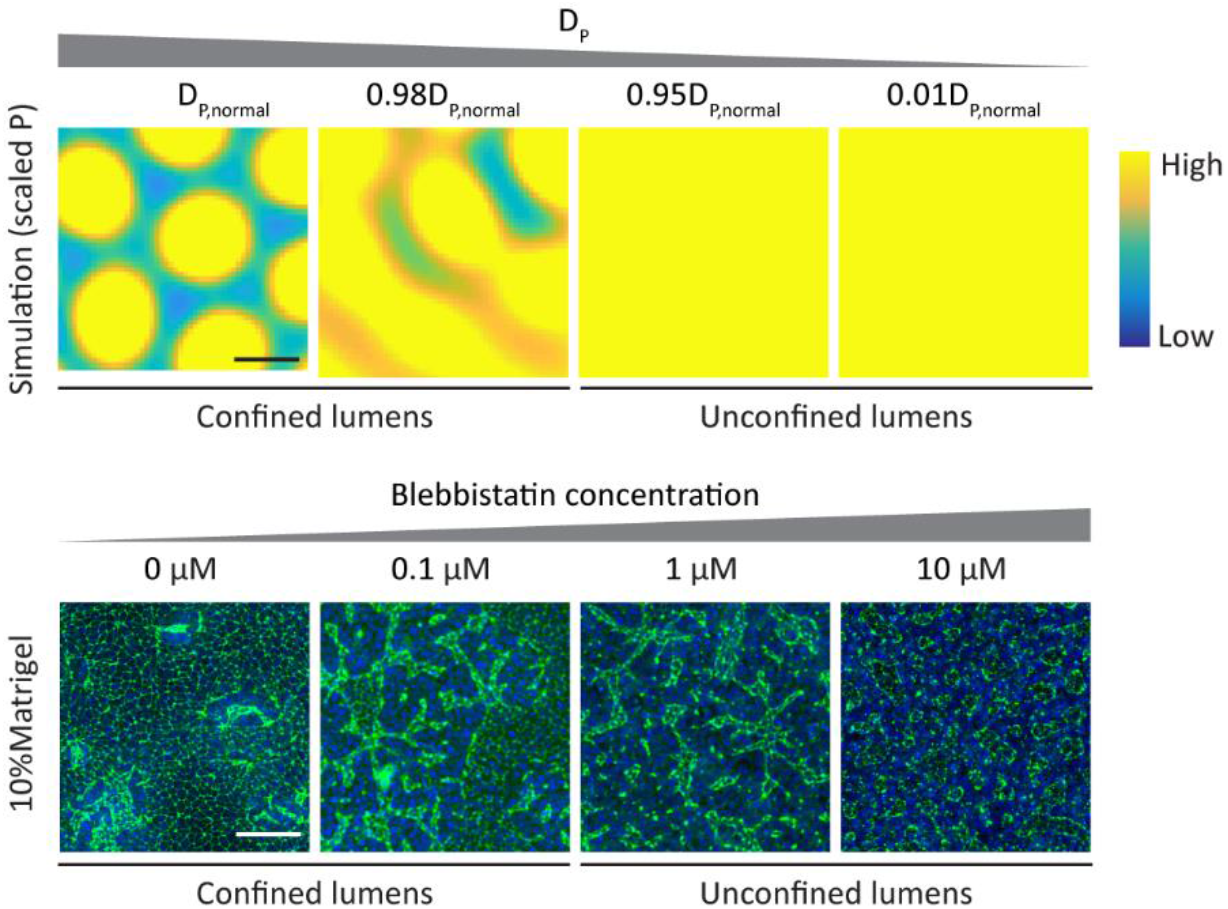
*In silico* and *in vitro* contractility perturbation comparison. Representative images of scaled P results under different values of D_p_. Representative images of ZO1 expression results under various Blebbistatin concentrations (n = 3).

## Materials and Methods

### Human PSC lines

The human PSC lines used in this study are 1) NCRM-1 (RRID:CVCL_1E71) hPSC line from NIH Center for Regenerative Medicine (CRM), Bethesda, USA. 2) ZO1 hPSC line (Mono-allelic mEGFP-Tagged TJP1 WTC, Coriell institute for Medical Research)

### Human PSC culture

Human PSC cultures were maintained in Matrigel-coated 6 well-plates until 60-70% confluence. To pass cells, a 4 m Dispase (Sigma-Aldrich) treatment was applied at 37°C by PBS washes. Colonies were gently scraped prior to gentle agitation to break down the colonies. The cell concentration was diluted 1:6 and cultured in 2 mL of Essential 8 (E8) - Flex Medium Kit (ThermoFisher Scientific) supplemented with 1% Penicillin Streptomycin (GIBCO) and Y-27632 ROCK inhibitor (ROCKi) (Hellobio) at 10 μM for 24 h. The medium was replaced with 4 mL of fresh ROCKi-free E8-Flex medium for an additional 48 h when cells reached 60-70% confluence.

### Matrigel experiments

Human hPSCs, at a confluence of 60–70%, were washed with PBS three times, followed by the application of 1 mL of TrypLE Express (GIBCO) at 37°C for 3 m for dissociation. Once dissociated into single cells, 9 mL of DMEM/F12 medium supplemented with 20% FBS (GIBCO) was added for neutralization and TrypLE Express washing. The cells were then centrifuged at 300 RCF for 3 m. After discarding the supernatant, a second wash was applied by adding 10 mL of DMEM/F12 medium containing 20% FBS followed by centrifugation at 300 RCF for 3 m. The supernatant was discarded and the cells were resuspended in 1 mL E8-Flex medium supplemented with 10 μM of ROCKi. A cell count was performed and a volume was added containing the appropriate amount of cells to cell-free medium (E8-Flex - 10 μM ROCKi) to obtain a cell density of 150,000 cells/mL. This amounts to 30,000 cell for 200 μL of media added to a 96 well-plate. The cells were left to incubate for 24 h. Next, the medium was changed fully with fresh ROCKi-free E8-Flex medium and the cell were left to incubate for an additional 24 h. At 48 h, with confluent wells, the cells are ready for Matrigel introduction. Matrigel was prepared and thawed on ice. Depending on the desired Matrigel concentration, pre-cooled (on ice) ROCKi-free E8-Flex medium was supplemented with 10%, 8%, 6%, 4%, 2%, 1% and 0.1% Matrigel. After properly mixing the Matrigel, 200 μL of the medium was added immediately to the cells and was kept for 24 h or 48 h. In all experiments Matrigel was introduced to the cells after 48 h of seeding. Control conditions also receive pre-cooled ROCKi-free E8-Flex medium to minimize confounding factors.

Differentiation experiments using Matrigel utilized mesoderm differentiation medium. The mesoderm differentiation medium was comprised of DMEM/F12 (GIBCO), 1% N2 (GIBCO), 2% B-27 (GIBCO), 1 mM sodium pyruvate MEM (GIBCO), 1 mM glutamax (GIBCO), 1 mM non-essential amino acids (GIBCO) and 2% Penicillin Streptomycin (GIBCO), 10 μM of CHIR99021 (Tocris) and 500 nM of LDN193189 (Stemgent), and 20 ng/mL of fibroblast growth factor 2 (FGF2) (Peprotech). At the Matrigel addition timepoint (48 h), the mesoderm medium was supplemented with Matrigel as desired and as previously described and then added to cells for an additional 48 h. The cells were then fixed, stained with mesoderm markers, imaged and analyzed. Control conditions did not received Matrigel.

For inhibition experiments, 10 μM Blebbistatin (Sigma–Aldrich), or 5 μM of EZRIN inhibitor NSC668394 (Sigma–Aldrich) was supplemented to E8-Flex medium and introduced to cells after a full medium change at day 1 for a 24 h pretreatment prior to Matrigel addition. After 24 h, similar inhibition media were created, supplemented with 10% Matrigel and introduced to the cells for an additional 24 h. Control conditions did not received Matrigel.

### Laminin labeling and live tracking

To fluorescently tag laminin present in Matrigel, while on ice, a 10% Matrigel E8-Flex medium was created. Anti-Laminin antibody (Invitrogen, PA1-16730) was added to a final concentrated of 1:500 and incubated on ice for 5 min. Next donkey-anti mouse Alexa Fluor 647 was added to the medium to a final concentration of 1:1000 and incubated on ice for 5 min (Invitrogen). 200 μL of the medium was then introduced to the cells cultured in a well of a 96 well-plate and incubated for 5 min at 37°C and 5% CO_2_. Confocal microscopy immediately followed to track laminin dynamics and ZO1 expression for 24 h.

### Fluorescent particles Matrigel tagging and live tracking

While on ice, a 10% Matrigel E8-Flex medium was created. The medium was supplemented with fluorescent particles (FluoSpheres Carboxylate-Modified Microspheres, 200 nm, Invitrogen) to a final concentration of 1:500 v/v. The medium was then introduced to the cells and incubated for 5 min at 37°C and 5% CO_2_. Confocal microscopy immediately followed to track bead displacements and ZO1 expression for 24 h or 48 h.

### Shaking experiment

150,000 cell were seeded into a plate of a 12 well-plate following standard Matrigel experiment protocol for media changes. At day 2, a 2% Matrigel E8-Flex medium tagged with fluorescent particles was introduced to the cells, the plate was immediately placed on an orbital shaker at 40 RPM and incubated for 5 min at 37°C and 5% CO_2_ for 24 h. Control Matrigel conditions were not placed on the shaker. Matrigel free conditions were present in both conditions.

### Polyethylene glycol hydrogel Experiments

Nondegradable polyethylene glycol (PEG) hydrogel was prepared with a 0.7 kPa stiffness as previously described^9^ and supplemented with 1:500 Fluorescent particles. Here, the cell volume fraction was replaced by milliQ water. 50 μL of the unpolymerized hydrogel was introduced to the cells, instead of Matrigel, and left for 20 m at room temperature to complete the polymerization process. After which 200 μL of media was added. To supplement the hydrogel with Matrigel, the same hydrogel premixture was used and Matrigel was added to a final concentration of 10% and replaced the same volume of MilliQ water. The combined PEG-Matrigel mixture was added to the cells and left for 20 m at room temperature to complete the polymerization process. Confocal microscopy immediately followed to track the bead displacement and ZO1 expression.

### Fluorescently labeled Iron oxide clusters (FCs)

50 mg of Fe_3_O_4_ nanoparticles (Sigma-Aldrich, 20-40 nm) were first weighed under sterile conditions and added to 1 mL of PBS media, in a 1.5 mL Eppendorf tube. Due to the lack of a surfactant, the particles cluster together forming ~1-2 μm sized clusters with a concentration of 50 mg/mL. The solution was pipetted thoroughly and subsequently sonicated for 5 m to break apart any aggregation. 2 μL of the suspended clusters is put into 1 mL of E8-Flex medium. Next fluorescent particles (FluoSpheres Carboxylate-Modified Microspheres, 200 nm, Invitrogen) were added to a final concentration of 1:5000 v/v and the solution sonicated for 2 min. The FP tagged the iron oxide clusters forming fluorescent clusters (FCs).

To label the surface of monolayer, immediately after sonication, 100 μL of the FCs were added to each well of a 96 well-plate and incubated for 5 min at 37°C and 5% CO_2_. Next the medium was discarded and replaced with E8-Flex medium supplemented with Matrigel at the desired concentration. Control conditions received Matrigel-free medium.

### Microscopy

A Nikon Ti2 Eclipse inverted microscope with incubation capability was used to image FCs dynamics in various Matrigel conditions and in control conditions using a ×10 air objective with 15 min imaging interval for 24 h. Confocal representative images were obtained using a Leica SP8 DIVE (Leica Microsystems) confocal microscope using a ×10 air objective or a ×20 water-emersion objective. The same microscope was with incubation and resonance scanner capability to capture FP and laminin dynamics using a using a ×10 air objective with 15 min imaging interval for 24 or 48 h. To capture actin processes and dynamics the same microscope was used with ×64 oil-emersion objective with 20 s imaging interval. Low magnification imaging was done using an EVOS (Invitrogen) inverted microscope using a ×4 air objective.

### Scanning electron microscopy (SEM)

The media compositions and exposure times followed the same protocol as the pluripotent conditions. First, hPSCs were grown on Matrigel-coated 13mm inside wells of a 24 well-plate for 24 h under ROCKi conditions, then an additional 24 h under ROCKi-free conditions. At the Matrigel addition timepoint, the Matrigel conditions received E8-Flex medium supplemented with 10% Matrigel and the controls remained Matrigel-free. All media volumes were adjusted to 500 μL due to the well size of the 24 well-plate. After 24 h of Matrigel exposure, cells were fixed with 2.5% glutaraldehyde in 0.1 M Na-cacodylate buffer Ph 7.2. After 3x washing in the same buffer, the coverslips with cells were incubated in 2% osmium tetroxide (Electron Microscopy Sciences, Hatfield PA, US) for 2 h, washed in milli-Q water and dehydrated in a graded ethanol series to 100% ethanol. Finally, keeping the coverslips submerged in 100% ethanol at all times, they were inserted in a coverslip-holder for critical point drying in a Leica CPD300 apparatus (Leica, Vienna, AT). Finally, the dried coverslips were mounted on 25mm SEM support stubs with Carbon-stickers, the edges were silver painted, and the assembly was coated with 10 nm Chromium in a Leica ACE600 coating machine (Leica, Vienna, AT). Cells were studied and imaged in a Zeiss Sigma SEM (Zeiss, Oberkochen, DE) at an accelerating voltage of 2kV and a working distance of 7mm.

### Particle image velocimetry

PIVlab^24,25^, was used in combination with MATLAB (MATLAB, R2018b, The MathWorks Inc.). Here FP time lapse images at the Matrigel-monolayer interface were uploaded to PIVlab through graphical user interface. Standard parameter settings were used along with the PIV FFT window deformation algorithm. A 3 pass analysis was implemented and a 128, 64 and 32 pixels interrogation areas were used for Pass 1, 2 and 3 respectively. Calibration was set and later verified using image scale to ensure correct velocity reporting. Data visualization used data smoothing at maximum strength and the standard strength for the high-pass vector field filter. This was visually verified to ensure vectors did not misreport velocity orientation and magnitude. Text data extracting instantaneous and average velocities were obtained. These text matrices were input into MATLAB to plot the streamlines for the average velocities and vectors for the instantaneous velocities on a contour map.

### Image analysis

ImageJ (Version 1.52a) was used for all image analysis. To obtain the velocity dynamics, gray scale velocity maps at 0, 6, 12, 18 and 24 h were extracted from PIVlab and input into ImageJ (Version 1.52a). Radial intensity profiles were obtained using the radial profile angle plugin around domes. The intensities were evaluated for the various time points. 5 velocity zones were radially marked and the velocities within averaged to obtain 5 velocity values for each time point.

To visualize the trajectories of FCs, 100 FCs per condition were manually tracked using the MTrackJ plugin on ImageJ. Using the data extracted from the trajectory analysis, average trajectory velocities, travel lengths and start to finish distances were calculated for each trajectory in different conditions.

Using the OrientationJ^26,27^ plugging, ZO1 orientation and coherence scores were obtained. Using the radial profile angle plugin around domes, radial profiles of ZO1 intensity, FC intensity and coherence scores were measured for each image in the sequence of images comprising a 24 h time-lapse. Values from different domes were combined, averaged and compiled into kymographs for ZO1 intensity, FC intensity and coherence scores.

To evaluate ZO1 polarization ahead and behind FCs, line sections where used to trace particle movement on a stack of images. ZO1 and FC kymographs were then evaluated and superimposed. Next, a 25 μm line section for each time increment was centered on the particle position in the kymographs and ZO1 and FC intensities were then evaluated. Here, the combined ZO1 intensity form various particles and at various time points in the +12.5 μm ahead of the particle position was compared to that in the −12.5 μm space behind the particle.

Inter-dome distance evaluation was done by marking the center of individual domes. The coordinates of each dome were then used to compute the distance to every other dome within the same field of view. For each dome only the shortest distance is reported, i.e. the nearest dome. Duplicate values were excluded from the data. A similar technique was used for *in silico* dome measurement, however here *in silico* coordinates where compared to the *in vitro* coordinates and nearest dome x-coordinate rank was reported against the *in silico* x-coordinate rank. The average difference in distance between *in silico* and *in vitro* nearest domes was then evaluated and reported.

### Reaction-diffusion simulation

A reaction-diffusion (RD) system was built based on the following Brusselator equations for the signals E and P,

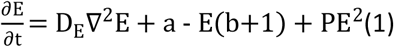

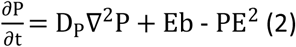

The parameters a and b control patterning outcomes from domes to stripes. We chose values of 5 and 7.75 for a and b respectively, which produced periodic domes in continuous domain spaces, and therefore reflecting *in vitro* observations. The E and P diffusivities were set for D_E_ = 20 and D_p_ = 160 respectively which reflected measures *in vitro* inter-dome distances under normal conditions. The diffusivity D_E_ was changed accordingly to reflected Blebbistatin treatment. MATLAB was used to solve a set of Laplacian operations in an iterative process. A spatial increment of 1 unit (10 μm) in both x and y-axis, a timestep of 0.001 s and a total time of 300 s were chosen as they provided a stable simulation. The domain space is by default continuous but can be rendered bounded in any direction by forcing desired edge values to 0 in every iteration. The domain space can be defined as matrix of m rows and n columns but can also be of any arbitrary shape by providing MATLAB a text matrix of 0s, unused spaces, and 1s used domain spaces. This binary matrix can also be obtained through a tissue mask created in ImageJ using the a saturated Hoescht signal from *in vitro* data. In this way, any tissue shape can be simulated in the RD model.

### Immunohistochemistry, actin labeling and patterning studies

After 24 or 48 h cells were fixed using 4% paraformaldehyde (Sigma–Aldrich) for 2 h followed by washing with PBS. Sample permeabilization and blocking was made using solution made with 0.3% Triton X (PanREAC AppliChem) and 0.5% BSA (Sigma–Aldrich) for 30 min. Primary antibodies were suspended in permeabilization and blocking solution and applied to the sample for 24 h to ensure diffusion through Matrigel. This was followed by three PBS washing over 24 h. Next immunolabeling was performed over a duration of 24 h using donkey-anti mouse Alexa Fluor 555/647 (Invitrogen, dilution 1:500), donkey-anti rabbit Alexa Fluor 555/647 (Invitrogen, dilution 1:500), or donkey-anti goat Alexa Fluor 555/647 (Invitrogen, dilution 1:500) secondary antibodies. To visualize filamentous actin Alexa Fluor 647 conjugated phalloidin was used (Invitrogen, dilution 1:4000). Hoechst was used to visualize nuclei. The ZO1 reported cell line was used to visualize ZO1 expression.

Actin was labeled to visualize live actin dynamics. At 24 h after Matrigel introduction, 100 μL of E8-Flex supplemented with 1 nM of SiR-actin (Spirochrom) was added to the cells and incubated for 10 min. Confocal microscopy was then conducted to observe apical actin dynamics.

The expression of NANOG was used to evaluate regions of high pluripotency in Matrigel and control cases. NANOG and T were used to evaluate to contrast mesoderm expression in differentiation experiments. Visual inspection of localization was conducted and reported.

### Single-cell RNA sequencing sample preparation

Samples were grown following the protocol for control and 10% Matrigel conditions in 6 well plates, 3 wells per condition. After 24 h of Matrigel exposure, samples were washed with PBS to ensure little to no Matrigel is left, then dissociated into single cells using an incubation period of 3 min in 1 mL TrypLE Express in every well. The 1 mL solutions were neutralized with 9 mL of DMEM medium supplemented with 20% FBS, the control and Matrigel conditions were pooled, then centrifuged for 5 min at 300 RCF. After decanting the supernatant, the cell pellets were resuspended in 200 μL of E8-Flex medium and put on ice. Library preparations for the scRNAseq was performed using 10X Genomics Chromium Single Cell 3’ Kit, v3 (10X Genomics, Pleasanton, CA, USA). The cell count and the viability of the samples were assessed using LUNA dual fluorescence cell counter (Logos Biosystems) and a targeted cell recovery of 10,000 cells was aimed for all the samples. Post cell count and QC, the samples were immediately loaded onto the HyDrop Hybrid^28^ platform. Single-cell RNA-seq libraries were prepared using manufacturers recommendations (Single-cell 3’ reagent kits v3 user guide; CG00052 Rev B), and at the different check points, the library quality was assessed using Qubit (ThermoFisher) and Bioanalyzer (Agilent). Sequencing was performed using BGI Genomics. The data generated from sequencing was demultiplexed and mapped against human genome reference using CellRanger v3.0.2.

### Single-cell sequencing data processing and clustering

After QC steps we have retained 6503 and 6746 sequenced cells for the control and 10% Matrigel conditions respectively. We implemented the Seurat^29^ 4.2.0 pipeline to scRNAseq analysis. Data normalization was performed, followed by the identification of 2000 highly variable genes using the FindVariableFeatures. Auto scaling of the data was performed and described using principal component analysis (PCA) using the RunPCA function. Data clustering was done using the top 15 PCs and the FindNeighbors function parameter k.param = 20. The FindCluster resolution was set to 0.5. Data visualization was done using the Uniform Manifold Approximation and Projection (UMAP) dimensionality reduction technique using the RunUMAP package with the top 15 PCs. Cluster annotation was performed primarily based on the top 25 genes and/or specific marker genes, which identified seven clusters, 1) Proliferation, glycolysis, 2) Non-proliferative, Glycolysis, 3) Non-proliferative, 4) Cycling, 5) Transition 1, 6) Transition 2, 7) Neural. Genesets were used to highlight certain identified processes, Microvilli (*PLS1*, *TWF2*, *VIL1*, *KLF5*, *PLD1*, *FSCN1*), Pluripotency (*NANOG*, *POU5F1*, *CDH1*), Proliferation (*TOP2A*, *MKI67*, *CDC42*), Cytoskeleton remodeling (*LIMK1*, *TJP1*, *ROCK1*, *ROCK2*, *MYH9*, *VCL*, *ACTA1*), ECM remodeling (*ITGB1*, *ITGB2*, *LAMA1*, *MMP9*), Glycolysis (*ALDOA*, *ENO1*, *ENO2*, *HK1*, *PFKL*, *PFKM*). Similarly, genesets were created for Epiblast, Primitive streak, and Nascent mesoderm identities following the top 25 markers of each of these clusters in the human gastrula dataset^30^.

### Differential gene expression analysis (DGEA), gene ontology (GO) enrichment analysis

We perform DGEA using the FindMarkers function in the Seurat R packages between 10% Matrigel and control cells with function parameters min.pct = 0.0 and logfc.threshold = 0.0.

The resultant total gene list was then ranked in decreasing expression levels. GO enrichment analyses was then conducted using the GOrilla^31^ web portal using the Homo sapiens organism single ranked file running mode and p-value threshold of 10^−5^. The results were then passed to the REVIGO^32^ web portal to obtain a summarized GO term list. Default settings were used.

### Hierarchical clustering

Hierarchical clustering was performed using geneset scores for the various clusters and the R package heatmap.2.

### Single R analysis

The SingleR^33^ package was used to project the cell of the dataset generated here onto the annotated clusters of the human gastrula (E-MTAB-9388)^30^. The top-50 markers genes (log fold change ≥0.5, min.pct ≥0.5) of the clusters identified in human gastrula data set were used as reference.

### Quantification and statistical analysis

Two-way ANOVA statistical tests and unpaired two-tailed t-test with corrections were used where appropriate with a 95% confidence interval (GraphPad Prism 6, Version 6.01, GraphPad Software, Inc.). Statistical significance was considered for all comparisons with p < 0.05.

### Data availability

All raw sequencing data, and the combined processed and metadata files generated in this study will become available at GEO.

## Acknowledgements

This work was supported by the FWO grant G087018N and FWO postdoctoral fellowship 1217220N, Interreg Biomat-on-Chip grant and Vlaams-Brabant and Flemish Government co-financing, KULeuven grantsC14/17/111, C32/17/027 and IDN/20/007, and by an Allen Distinguished Investigator Award, a Paul G. Allen Frontiers Group advised grant of the Paul G. Allen Family Foundation.

